# PPM-Decay: A Computational Model of Auditory Prediction with Memory Decay

**DOI:** 10.1101/2020.01.09.900266

**Authors:** Peter M. C. Harrison, Roberta Bianco, Maria Chait, Marcus T. Pearce

**Author notes:** Author Note: This is an unpublished preprint that has yet to undergo peer review (January 10, 2020). Peter M. C. Harrison, School of Electronic Engineering and Computer Science, Queen Mary University of London; Roberta Bianco, Ear Institute, University College London; Maria Chait, Ear Institute, University College London; Marcus T. Pearce, School of Electronic Engineering and Computer Science, Queen Mary University of London. Peter Harrison is now at the Max Planck for Empirical Aesthetics, Frankfurt, Germany. He was previously supported by a doctoral studentship from the EPSRC and AHRC Centre for Doctoral Training in Media and Arts Technology (EP/L01632X/1). Correspondence concerning this article should be addressed to Peter M. C. Harrison, Max-Planck-Institut für empirische Ästhetik, Grüneburgweg 14, 60322 Frankfurt am Main, Germany.

## Abstract

Statistical learning and probabilistic prediction are fundamental processes in auditory cognition. A prominent computational model of these processes is Prediction by Partial Matching (PPM), a variable-order Markov model that learns by internalizing *n*-grams from training sequences. However, PPM has limitations as a cognitive model: in particular, it has a perfect memory that weights all historic observations equally, which is inconsistent with memory capacity constraints and recency effects observed in human cognition. We address these limitations with PPM-Decay, a new variant of PPM that introduces a customizable memory decay kernel. In three studies – one with artificially generated sequences, one with chord sequences from Western music, and one with new behavioral data from an auditory pattern detection experiment – we show how this decay kernel improves the model’s predictive performance for sequences whose underlying statistics change over time, and enables the model to capture effects of memory constraints on auditory pattern detection. The resulting model is available in our new open-source R package, *ppm* (https://github.com/pmcharrison/ppm).

---

Humans are sensitive to structural regularities in sound sequences (Agres, Abdallah, & Pearce, 2018; Barascud, Pearce, Griffiths, Friston, & Chait, 2016; Bendixen, Schroger, & Winkler, 2009; Cheung, Meyer, Friederici, & Koelsch, 2018; Garrido, Sahani, & Dolan, 2013; Koelsch, Busch, Jentschke, & Rohrmeier, 2016; Rohrmeier et al., 2012; Tillmann & Poulin-Charronnat, 2010; Wacongne et al., 2011; Winkler, Denham, & Nelken, 2009). This structural sensitivity underpins many aspects of audition, including sensory processing (Southwell & Chait, 2018; Turk-Browne, Scholl, Johnson, & Chun, 2010), auditory scene analysis (Andreou, Kashino, & Chait, 2011; Schröger et al., 2014), language acquisition (Erickson & Thiessen, 2015), and music perception (Pearce, 2018).

The Prediction by Partial Matching (PPM) algorithm is a powerful approach for modeling this sensitivity to sequential structure. PPM is a variable-order Markov model originally developed for data compression (Cleary & Witten, 1984) that predicts successive tokens in symbolic sequences on the basis of *n*-gram statistics learned from these sequences. An *n*-gram is a contiguous sequence of *n* symbols, such as “ABA” or “ABB”; an *n*-gram model generates conditional probabilities for symbols, for example the probability that the observed sequence “AB” will be followed by the symbol “A”, based on the frequencies of different *n*-grams in a training corpus. Different values of *n* yield different operational characteristics: in particular, small values of *n* are useful for generating reliable predictions when training data are limited, whereas large values of *n* are useful for generating more accurate predictions once sufficient training data have been obtained. The power of PPM comes from combining together multiple *n*-gram models with different orders (i.e. different values of *n*), with the weighting of these different orders varying according to the amount of training data available. This combination process allows PPM to retain reliable performance on small training datasets while outperforming standard Markov chain models with larger training datasets.

The PPM algorithm has been adopted by cognitive scientists and neuroscientists as a cognitive model for how human listeners process auditory sequences. The algorithm has proved particularly useful in modeling music perception, forming the basis of the Information Dynamics Of Music (IDyOM) model of Pearce (2005) which has been successfully applied to diverse musical phenomena such as melodic expectation (Pearce & Wiggins, 2006), emotional experience (Egermann, Pearce, Wiggins, & McAdams, 2013), similarity perception (Pearce & Müllensiefen, 2017), and boundary detection (Pearce, Müllensiefen, & Wiggins, 2010). More recently, the PPM algorithm has been applied to non-musical auditory modeling, including the acquisition of auditory artificial grammars (Agres et al., 2018) and the detection of repeating patterns in fast tone sequences (Barascud et al., 2016).

These cognitive studies typically use PPM as an *ideal-* or *rational-observer* model. Applied to a particular experimental paradigm, an ideal-observer model simulates a theoretically optimal strategy for performing the participant’s task. This optimal strategy provides a benchmark against which human performance can be measured; deviations from this benchmark can then be analysed to yield further insights into human cognition. In artificial experimental paradigms, where the stimuli are generated according to a prespecified formal model, it is often possible to derive a “true” ideal-observer model that provably attains optimal performance. However, in naturalistic domains (e.g. music, language) the researcher does not typically have access to the true model that generated the stimuli, and so it is not possible to construct a provably optimal ideal-observer model. Moreover, in certain experimental paradigms (e.g. fast auditory pattern detection, Barascud et al., 2016) it is unlikely that the participant’s cognitive processes reflect a strategy perfectly tailored to the exact experimental task; instead, they are likely to reflect general principles that tend to work well for naturalistic perception. PPM is typically applied in these latter contexts: it does not constitute the provably optimal observer for most particular tasks, but it represents a rational model of predictive processing that is assumed to approximate ideal performance for a broad variety of sequential stimuli.

However, the PPM algorithm suffers from an important limitation when applied to cognitive modeling. All observed data are stored in a single homogenous memory unit, with historic observations receiving equal salience to recent observations. This is problematic for two reasons. First, it means that the model performs suboptimally on sequences where the underlying statistical distribution changes over time. Second, it means that the model cannot capture how human memory separates into distinct stages with different capacity limitations and temporal profiles, and the way that these different stages interact to determine cognitive performance (e.g. Atkinson & Shiffrin, 1968; Nees, 2016; Neisser, 1967). While various sequence modeling approaches from the cognitive literature do incorporate phenomena such as recency effects and capacity limits (Bröker, Bestmann, Dayan, & Marshall, 2018; Harrison, 2011; Mattar, Kahn, Thompson-Schill, & Aguirre, 2016; Meyniel, Maheu, & Dehaene, 2016; Norton, Fleming, Daw, & Landy, 2017; O’Reilly, 2013; Skerritt-Davis & Elhilali, 2018, 2019; Squires, Wickens, Squires, & Donchin, 1976; Yu & Cohen, 2008) these approaches are generally limited to low-order statistics and cannot therefore match the predictive power of PPM. Conversely, more powerful sequence models from the machine-learning literature are difficult to tailor to the idiosyncrasies of human memory (e.g. hidden Markov models, Rabiner, 1989; long short-term memory recurrent neural networks, Hochreiter & Schmidhuber, 1997).

Several partial solutions to this problem have been presented in the PPM literature. Moffat’s (1990) implementation allocated a fixed amount of storage space to the trie data structure used to store observed data, and rebuilt this tree from scratch each time this storage limit was exceeded, after Cormack & Horspool (1986). This solution may be computationally efficient but it has limited cognitive validity. Conklin & Witten (1995) introduced a technique whereby two PPM models would be trained, a long-term model and a short-term model, with the long-term model retaining training data from all historic sequences and the short-term model only retaining training data from the current sequence. The predictions from these two models would then be combined to form one probability distribution. This technique works well for capturing the distinction between the structural regularities characterizing a domain (e.g. a musical style, a language) and the statistical regularities local to a given item from the domain (e.g. a musical composition or a specific text), but it cannot capture recency effects within a given sequence or distinguish between historic sequences of different vintages.

Here we present a new version of the PPM algorithm that directly addresses these issues of memory modeling. This new algorithm, termed “PPM-Decay”, introduces a decay kernel that determines the weighting of historic data as a function of various parameters, typically the time elapsed since the historic observation, or the number of subsequent observations (Figure 1). It also introduces stochastic noise into memory retrieval, allowing the model to capture analogous imperfections in human memory. We have developed an open-source implementation of the model in C++, made available in the R package *ppm*, that allows the user to configure and evaluate different variants of the PPM-Decay model on arbitrary sequences.

**Figure 1 .**
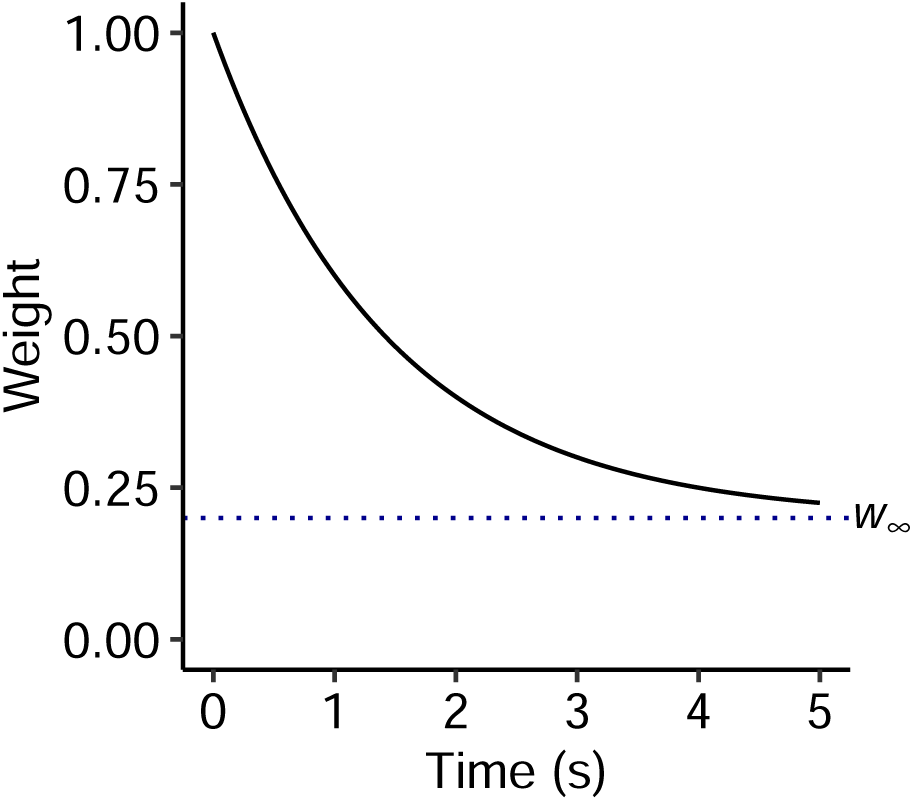
A simple decay kernel with an initial weight *w*_0_ = 1, an exponential decay with half life *t*_0.5_ = 1 s, and an asymptotic weight *w*_∞_ = 0.2.

We demonstrate the utility of this new algorithm in a series of experiments corresponding to a variety of task domains. Experiment 1 simulates the prediction of sequences generated from a prespecified statistical model, and shows that incorporating memory decay improves the predictive performance of PPM for sequences when the underlying model parameters change over time. Experiment 2 simulates the prediction of chord sequences from three musical styles, and shows that a decay profile with a non-zero asymptote is useful for capturing a combination of statistical regularities specific to the current composition and statistical regularities general to the musical style. Experiment 3 models new empirical data from human listeners instructed to detect repeated patterns in fast tone sequences, and shows that a multi-stage decay kernel is useful for explaining human performance. Together these experiments speak to the utility of the PPM-Decay algorithm as a cognitive model of symbolic sequence processing.

## Experiment 1: Memory decay helps predict sequences with changing statistical structure

The original PPM algorithm weights all historic observations equally when predicting the next symbol in a sequence. This represents an implicit assertion that all historic observations are equally representative of the sequence’s underlying statistical model. However, if the sequence’s underlying statistical model changes over time, then older observations will be less representative of the current statistical model than more recent observations. In such scenarios, an ideal observer should allocate more weight to recent observations than historic observations when predicting the next symbol.

Various weighting strategies can be envisaged representing different inductive biases about the evolution of the sequence’s underlying statistical model. A useful starting point is an exponential weighting strategy, whereby an observation’s salience decreases by a constant fraction every time step. Such a strategy is biologically plausible in that the system does not need to store a separate trace for each historic observation, but instead can simply maintain one trace for each statistical regularity being monitored (e.g. one trace per *n*-gram), which is incremented each time the statistical regularity is observed and decremented automatically over time. This exponential-weighting strategy can also be rationalised as an approximation to optimal Bayesian weighting for certain types of sequence structures (Yu & Cohen, 2008).

We will now describe a proof-of-concept experiment to demonstrate the intuitive notion that such weighting strategies can improve predictive performance in the PPM algorithm. This experiment used artificial symbolic sequences generated from an alphabet of five symbols, where the underlying statistical model at any particular point in time was defined by a first-order Markov chain. A first-order Markov chain defines the probability of observing each possible symbol conditioned on the immediately preceding symbol; second-order Markov chains are Markov chains that take into account two preceding symbols, whereas zeroth-order Markov chains take into account zero preceding symbols. Our sequence-generation models were designed as hybrids between zeroth-order and first-order Markov chains, reflecting PPM’s capacity to model sequential structure at different Markov orders. These generative models took the form of first-order Markov chains, where each first-order conditional distribution was sampled from a symmetric Dirichlet prior with concentration parameter 0.1, and then averaged with a common zeroth-order distribution sampled from the same Dirichlet prior. These models can be represented as two-dimensional transition matrices, where the cell in the *i*th row and the *j*th column identifies the probability of observing symbol *j* given that the previous symbol was *i* (Figure 2A). Zeroth-order structure is then manifested as correlations between transition probabilities in the same column, and can be summarised in marginal bar plots (Figure 2A).

**Figure 2 .**
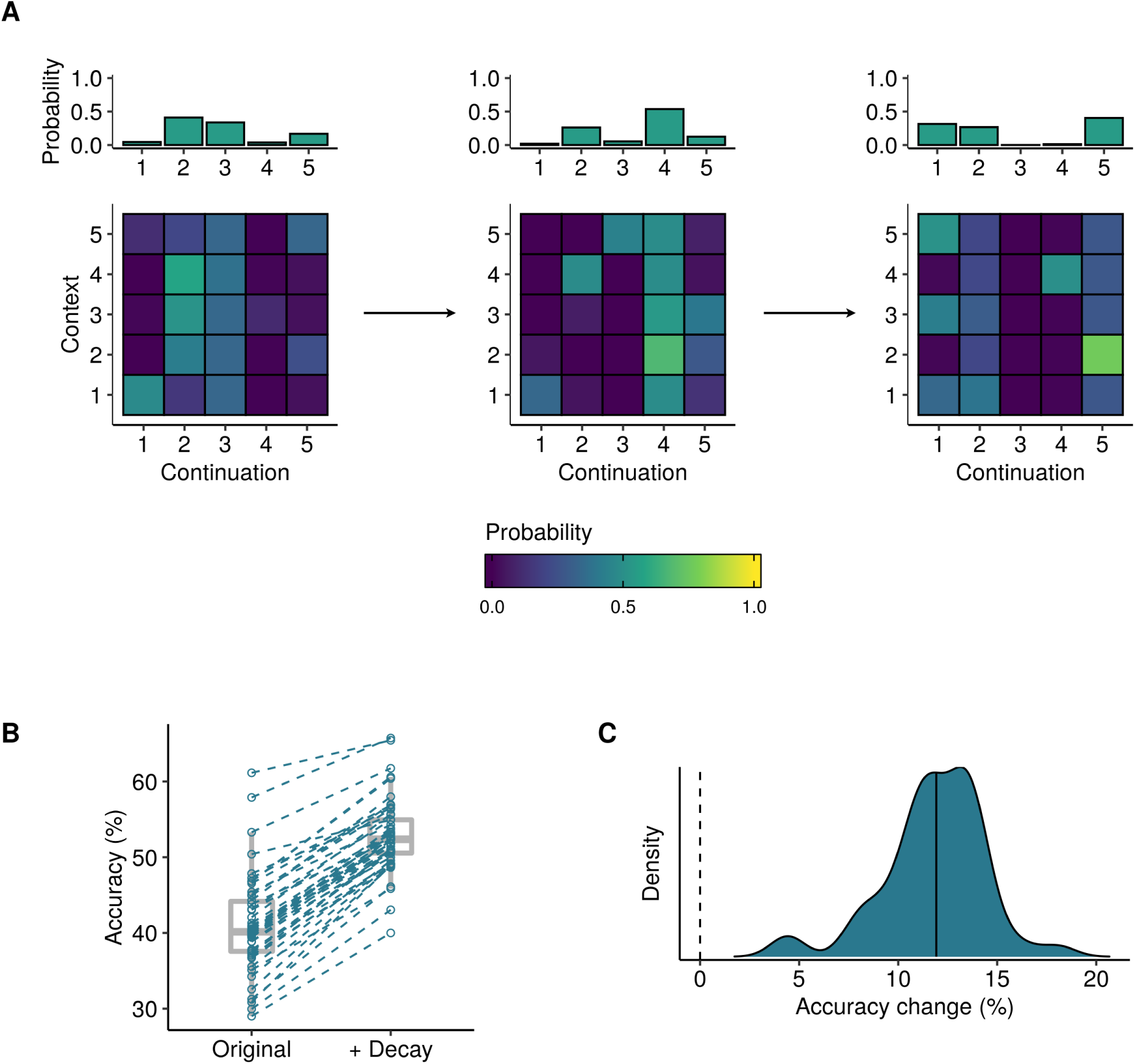
Illustrative plots for Experiment 1. **A)** Example sequence-generation models as randomly generated in Experiment 1. The bar plots describe 0th-order symbol distributions, whereas the matrices describe 1st-order transition probabilities. **B)** Repeated-measures plot indicating how predictive accuracy for individual sequences (*N* = 500, hollow circles) increases after the introduction of an exponential-decay kernel. **C)** Absolute changes in predictive accuracy for individual sequences, as summarised by a kernel density estimator. The median accuracy change is marked with a solid vertical line.

Each sequence began according to an underlying statistical model constructed by the above procedure, with the first symbol in each sequence being sampled from the model’s stationary distribution. At the next symbol, the underlying statistical model was either preserved with probability .99 or discarded and regenerated with probability .01. The new symbol was then sampled from the resulting statistical model conditioned on the immediately preceding symbol. This procedure was repeated to generate a sequence totalling 500 symbols in length.

Individual experimental trials were then conducted as follows. The PPM-Decay model was presented with one symbol at a time from a sequence constructed according to the procedure defined above, and instructed to return a predictive probability distribution for the next symbol. A single prediction was then extracted from this probability distribution, corresponding to the symbol assigned the highest probability. Prediction success was then operationalized as the proportion of observed symbols that were predicted correctly.

This experimental paradigm was used to evaluate a PPM-Decay model constructed with an exponential-decay kernel and a Markov order bound of one. This kernel is parametrized by a single half-life parameter, defined as the time interval for an observation’s weight to decrease by 50%. This half-life parameter was optimized by evaluating the model on 500 experimental trials generated by the procedure described above, maximizing mean prediction success over all trials using Rowan’s (1990) Subplex algorithm as implemented in the NLopt package (Johnson, 2019), and refreshing the model’s memory between each trial. The resulting half-life parameter was 12.26. The PPM-Decay model was then evaluated with this parameter on a new dataset of 500 experimental trials and compared with an analogous PPM model without the decay kernel.

The results are plotted in Figures 2B and 2C. They indicate that the exponential-decay kernel reliably improves the model’s performance, with the median percentage accuracy increasing from 48.8% to 62.2%. The exponential-decay kernel causes the algorithm to downweight historic observations, which are less likely to be representative of the current sequence statistics, thereby helping the algorithm to develop an effective model of the current sequence statistics and hence generate accurate predictions. Correspondingly, we can say that the exponential-decay model better resembles an ideal observer than the original PPM model.

## Experiment 2: Memory decay helps predict musical sequences

We now consider a more complex task domain: chord sequences in Western music. In particular, we imagine a listener who begins with zero knowledge of a musical style, but incrementally acquires such knowledge through the course of musical exposure, and uses this knowledge to predict successive chords in chord sequences. This process of musical prediction is thought to be integral to the aesthetic experience of music, and so it is of great interest to music theorists and psychologists to understand how these predictions are generated (Harrison & Pearce, 2018; Hedges & Wiggins, 2016; Pachet, 1999; Pearce, 2018; Rohrmeier & Graepel, 2012).

Chord sequences in Western music resemble sentences in natural language in the sense that they can be modeled as sets of symbols drawn from a finite dictionary and arranged in serial order. Such chord sequences provide the structural foundation of most Western music. For the purpose of modeling with the PPM algorithm, it is useful to translate these chord sequences into sequences of integers, which we do here using the mapping scheme described in *Methods*. For example, the first eight chords of the Bob Seger song “Think of Me” might be represented as the integer sequence “213, 159, 33, 159, 213, 159, 33, 159”.

Here we consider chord sequences drawn from three musical corpora: a corpus of popular music sampled from the Billboard “Hot 100” charts between 1958 and 1991 (Burgoyne, 2011), a corpus of jazz standards sampled from an online forum for jazz musicians (Broze & Shanahan, 2013), and a corpus of 370 chorale harmonizations by J. S. Bach (Sapp, 2005), translated into chord sequences using the chord labeling algorithm of Pardo and Birmingham (2002; see Methods for details). These three corpora may be taken as rough approximations of three musical styles: popular music, jazz music, and Bach chorale harmonizations. While we expect these three corpora each to be broadly consistent with general principles of Western tonal harmony (Piston, 1948), we also expect each corpus to possess distinctive statistical regularities that differentiate the harmonic languages of the three musical styles (Broze & Shanahan, 2013; Clercq & Temperley, 2011; Rohrmeier & Cross, 2008; Temperley & De Clercq, 2013). Figure 3 displays example chord sequences from these three corpora, alongside their corresponding integer encodings.

**Figure 3 .**
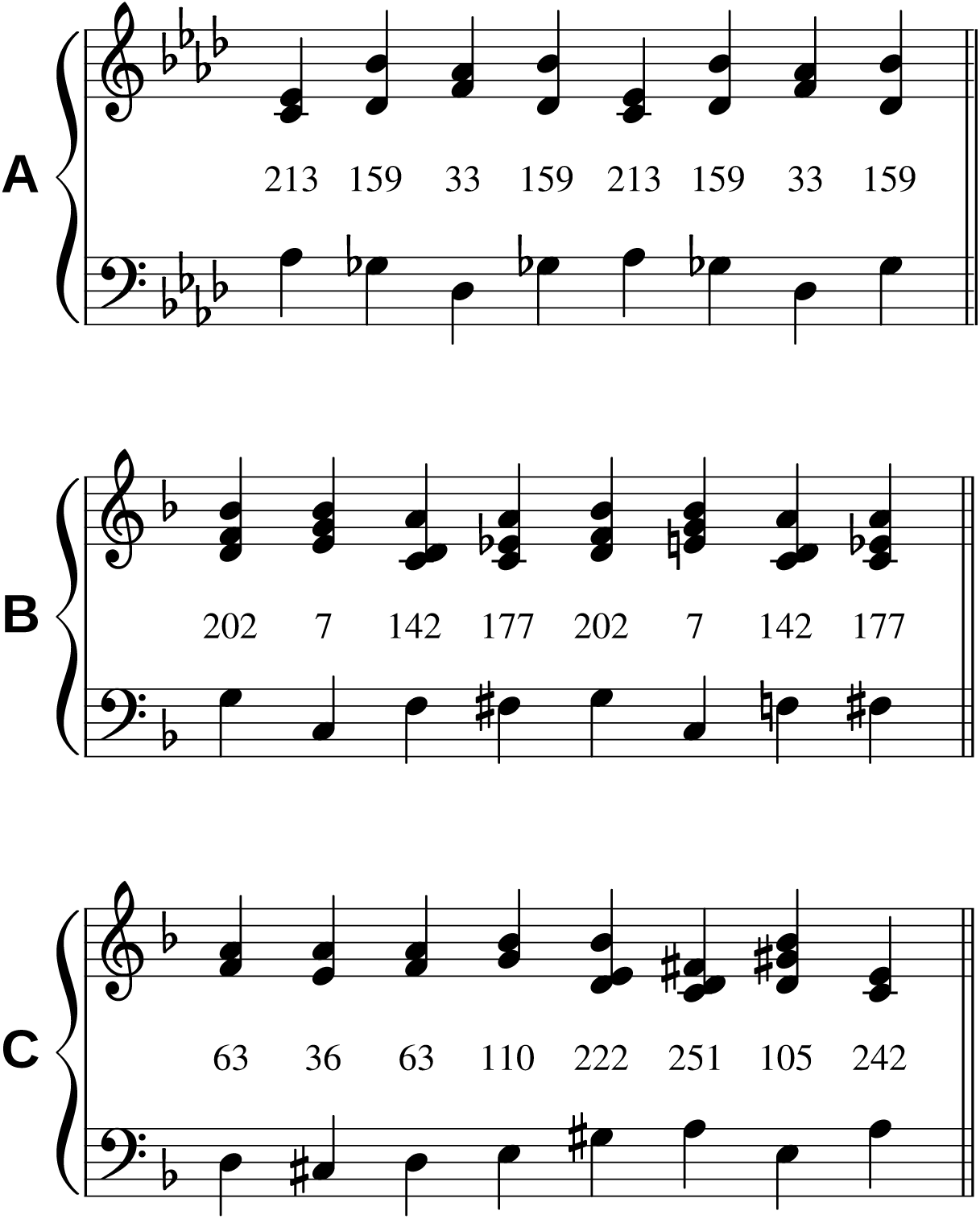
Sample chord sequences from **A)** the popular music corpus (“Night Moves”, by Bob Seger), **B)** the jazz corpus (“Thanks for the Memory”, by Leo Robin), and **C)** the Bach chorale harmonization corpus (“Mit Fried und Freud ich fahr dahin”, by J. S. Bach). Each chord is labeled by its integer encoding within the chord alphabet for the respective corpus. Each chord sequence corresponds to the first eight chords of the first composition in the downsampled corpus. Each chord is defined by a combination of a bass pitch class (lower stave) and a collection of non-bass pitch classes (upper stave). For visualization purposes, bass pitch classes are assigned to the octave below middle C, and non-bass pitch classes to the octave above middle C.

We expect the underlying sequence statistics to vary as we progress through a musical corpus. Sequence statistics are likely to change significantly at the boundaries between compositions, but they are also likely to change within compositions, as the chord sequences modulate to different musical keys, and travel through different musical sections. Similar to Experiment 1, we might therefore hypothesize that some kind of decay kernel should help the listener maintain an up-to-date model of the sequence statistics, and thereby improve predictive performance.

However, unlike Experiment 1, the chord sequences within a given musical corpus are likely to share certain statistical regularities. If the corpus is representative of a given musical style, then these statistical regularities will correspond to a notion of “harmonic syntax”, the underlying grammar that defines the harmonic conventions of that musical style. An ideal model will presumably take advantage of these stylistic conventions. However, the exponential-decay kernel from Experiment 1 is not well-suited to this task, because observations from historic sequences continuously decay in weight until they make essentially no contribution to the model. This is not ideal because these historic sequences will still contribute useful information about the musical style. Here we therefore evaluate a modified exponential-decay kernel, where memory traces decay not to zero but to a positive asymptote (see e.g. Figure 1). Such a kernel should provide a useful compromise between following the statistics of the current musical passage and capturing long-term knowledge of a style’s harmonic syntax.

We conducted our experiment as follows. For each musical corpus, we simulated a listener attempting to develop familiarity with the musical style by listening to one chord sequence every day, corresponding to one composition randomly selected from the corpus without repetition, for 100 days. We supposed that the listener began each chord sequence at the same time of day, so that the beginning of each successive chord sequence would be separated by 24-hour intervals, and we supposed that each chord in each chord sequence lasted one second in duration. Similar to Experiment 1, we supposed that the listener constantly tried to predict the next chord in the chord sequence, but this time we operationalized predictive success using the cross-entropy error metric, defined as the mean negative log probability of each chord symbol as predicted by the model. This metric is more appropriate than mean success rate for domains with large alphabet sizes, such as harmony, because it assigns partial credit when the model predicts the continuation with high but non-maximal probability. We used this metric to evaluate two decay kernels: the exponential-decay kernel evaluated in Experiment 1, termed the “Decay only” kernel, and a new exponential-decay kernel incorporating a positive asymptote, termed the “Decay + long-term learning” model. We found optimal parametrizations for these kernels using the same optimizer as Experiment 1 (the “Subplex” algorithm of Rowan, 1990), and compared the predictive performance of the resulting optimized models to a standard PPM model without a decay kernel. Each model was implemented with a Markov order bound of four, which seems to be a reasonable upper limit for the kinds of Markovian regularities present in Western tonal harmony (see e.g. Hedges, Roy, & Pachet, 2014; Landsnes et al., 2019; Rohrmeier & Graepel, 2012).

Figure 4 describes the performance of these two decay kernels. Examining the results for the three different datasets, we see that the utility of different decay parameters depends on the musical style. For the popular music corpus, incorporating exponential decay improves the model’s performance by c. 1.9 bits, indicating that individual compositions carry salient short-term regularities that the model can better leverage by downweighting historic observations. Introducing a non-zero asymptote to the decay kernel does not improve predictive performance on this dataset, indicating that long-term syntactic regularities contribute very little to predictive performance over and above these short-term regularities in popular music. A different pattern is observed for the jazz and Bach chorale corpora, however. In both cases, the decay-only model performs no better than the original PPM model, presumably because any improvement in capturing local statistics is penalized by a corresponding deterioration in long-term syntactic learning. However, incorporating a non-zero asymptote in the decay kernel allows the model both to upweight local statistics and still achieve long-term syntactic learning, thereby improving predictive performance by c. 1.5 bits.

**Figure 4 .**
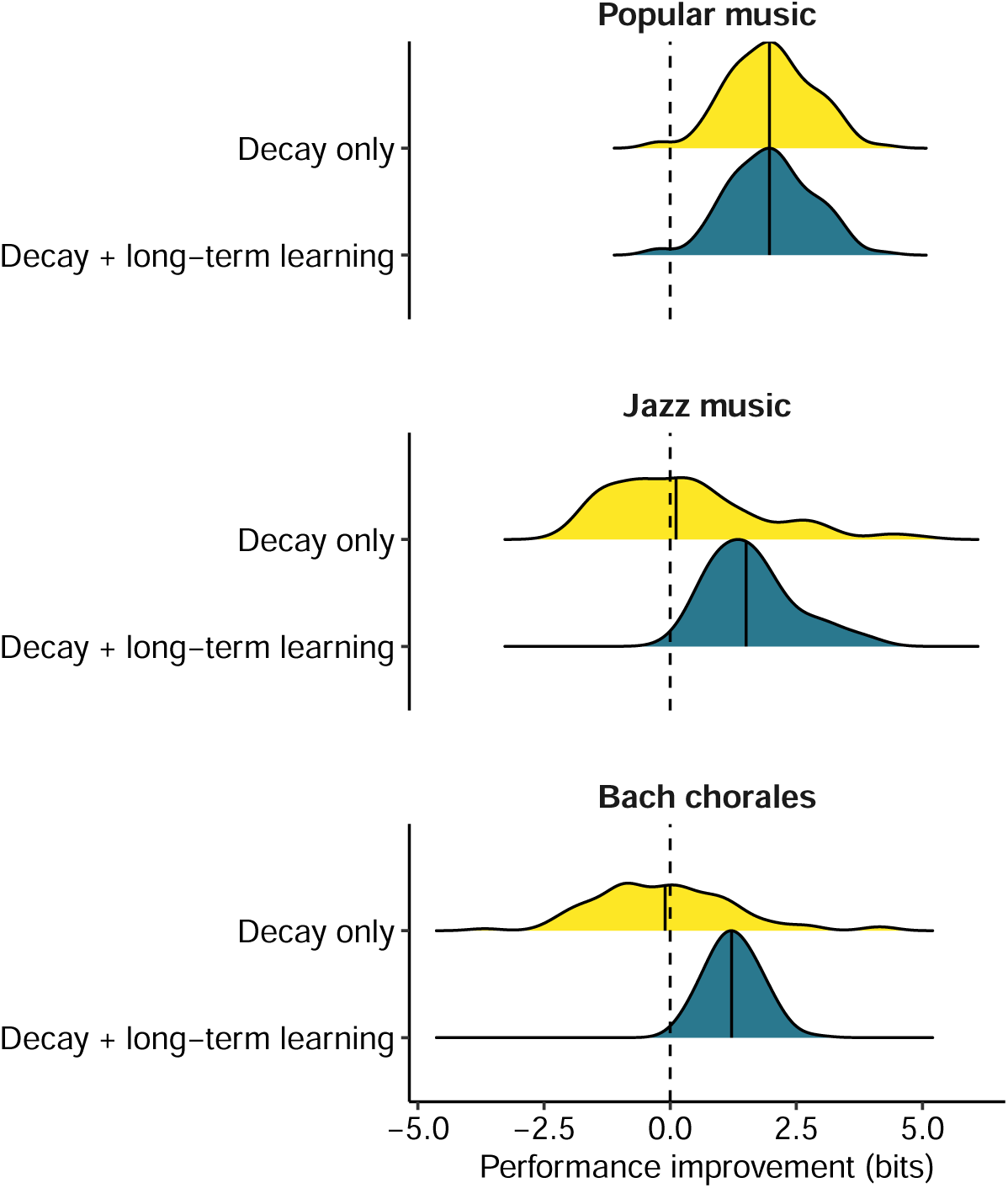
Predictive performances for different decay kernels in Experiment 2. Each composition contributed one cross-entropy value for each decay kernel; these cross-entropy values are expressed relative to the cross-entropy values of the original PPM model, and then summarised using kernel density estimators. Median performance improvements are marked with solid vertical lines.

These analyses have two main implications. First, they show that more advanced decay kernels are useful for producing a predictive model that better approximates ideal performance in the cognitive task of harmony prediction. The nature of these improved kernels can be related directly to the statistical structure of Western music, where compositions within a given musical style tend to be characterized by local statistical regularities, yet also share common statistical structure with other pieces in the musical style. An ideal-observer model of harmony prediction ought therefore to recognize these different kinds of statistical structure. Second, these analyses offer quantitative high-level insights into the statistical characteristics of the three musical styles. In particular, the popular music analyses found that long-term learning offered no improvement over a simple exponential-decay kernel, implying that the harmonic structure of popular music is dominated by local repetition. In contrast, both the jazz analyses and the Bach chorale analyses found that both exponential decay and long-term learning were necessary to improve from baseline performance, implying that chord progressions in these styles reflect both short-term statistics and long-term syntax to significant degrees.

## Experiment 3: Memory decay helps to explain the dynamics of auditory pattern detection

The PPM model has recently been used to simulate how humans detect recurring patterns in fast auditory sequences (Barascud et al., 2016). Barascud et al. used an experimental paradigm where participants were played fast tone sequences derived from a finite pool of tones, with the sequences organised into two sections: a random section (labelled “RAND”) and a regular section (labelled “REG”). The random section was constructed by randomly sampling tones from the frequency pool, whereas the regular section constituted a “frozen” sequence of frequencies from the pool which repeated identically for several iterations. These two-stage sequences, termed “RANDREG” sequences, were contrasted with “RAND” sequences which solely comprised one random section. The participant’s task was to detect transitions from random to regular sections as quickly as possible (see *Methods* for more details, and Figure 7 for an example trial).

**Figure 7 .**
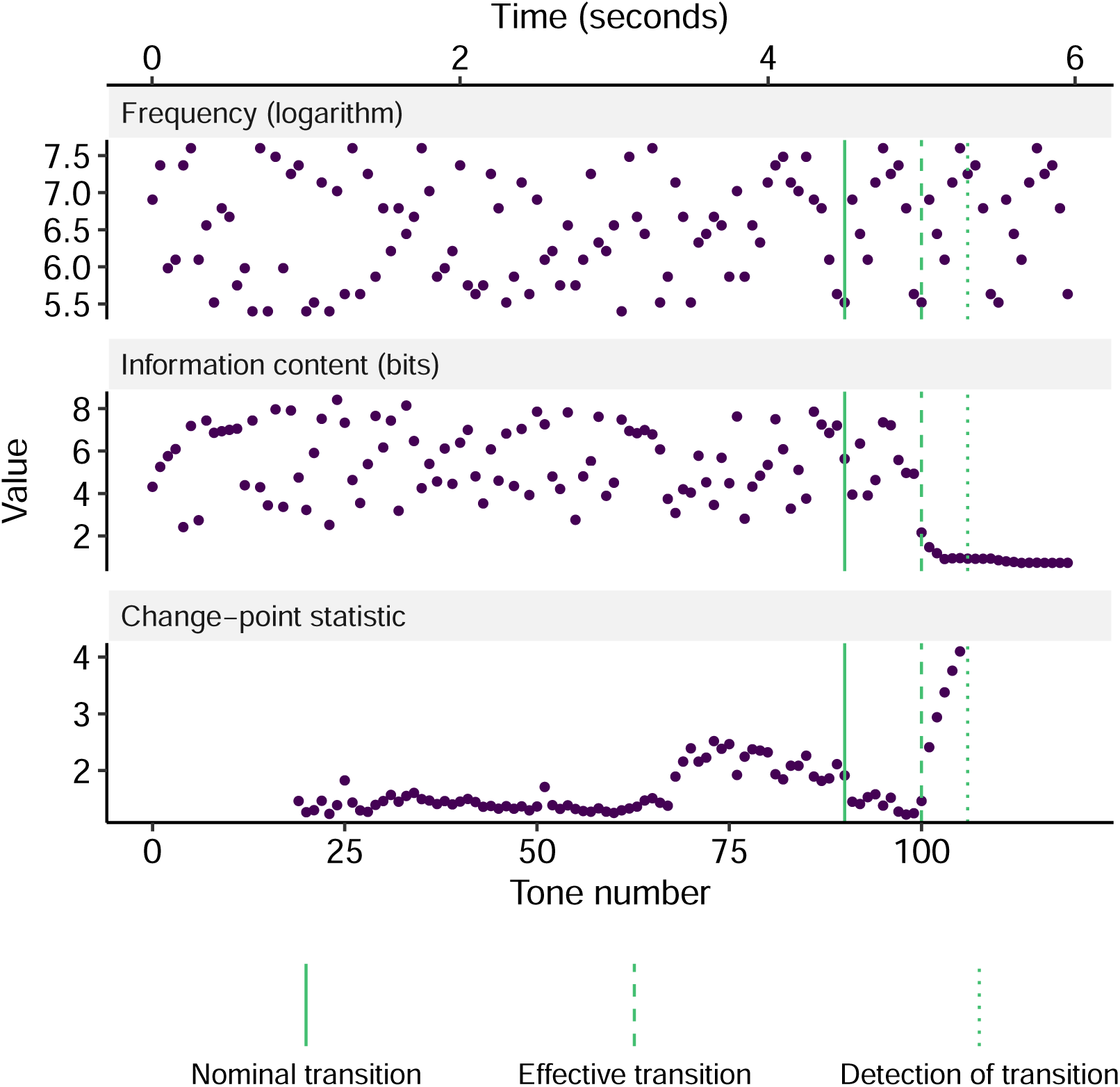
Example analysis of a single trial. The three panels plot each tone’s frequency, change-point statistic, and information content respectively. “Phase change” denotes the point at which the pattern changes from random tones to a repeating pattern of length 10. This repetition starts to become discernible after 10 tones (“First repetition”), at which point the sequence becomes fully deterministic. Correspondingly, information content (or “surprise”) drops, and triggers change-point detection at “Detection of transition”.

These experimental stimuli were constructed according to a well-defined statistical process, and it would be straightforward to derive a model that achieves provably optimal performance on the task given a well-defined performance metric. However, Barascud et al. reasoned that the cognitive mechanisms underlying fast auditory pattern recognition would be unlikely to be tailored to exact repetition, because exact repetitions are uncommon in naturalistic auditory environments. Instead, they supposed that human performance would be better characterized by more generic regularity detection mechanisms, such as those embodied in the PPM algorithm.

In particular, Barascud et al. (2016) suggested that listeners maintain an internal predictive model of incoming tone sequences that is incrementally updated throughout each sequence, and that listeners monitor the moment-to-moment surprise experienced by this model. They modeled this process using PPM as the predictive model, and operationalized surprise as the information content of each tone, defined as the tone’s negative log probability conditioned on the portion of the sequence heard so far. The authors proposed that listeners detect section changes based on the evolution of information content throughout the stimulus; in particular, changes from random to regular precipitate a sharp drop in information content, reflecting the transition from unpredictability to predictability.

Examining information content profiles produced by the PPM model, Barascud et al. (2016) concluded that an ideal observer should detect the transition from random to regular sections by the fourth tone of the second occurrence of the regular tone cycle. Analyzing behavioral and neuroimaging data, the authors found that participants reached this benchmark when the cycle length was small (5, 10 tones) but not when it was large (15, 20 tones). In other words, the ideal-observer model replicated human performance well for short cycle lengths, but some kind of cognitive constraints seemed to impair human performance for large cycle lengths.

One candidate explanation for this impaired performance is the limited capacity of auditory short-term memory. In order to detect a cycle repetition, the listener must compare incoming tones to tones that occurred at least one cycle ago. To achieve this, the listener’s auditory short-term memory must therefore span at least one cycle length. Short cycles may fit comfortably in the listener’s short-term memory, thereby supporting near-optimal task performance, but longer cycles may progressively test the limits of the listener’s memory capacity, resulting in progressively worsened performance.

An important question is whether this memory capacity is determined by temporal limits or informational limits. A temporal memory limit would correspond to a fixed time duration, within which events are recalled with high precision, and outside of which recall performance suffers. Analogously, an informational limit would correspond to a fixed number of tones that can be recalled with high fidelity from short-term memory, with attempts to store larger numbers of tones resulting in performance detriment.

Both kinds of capacity limits have been identified for various stages of auditory memory. Auditory sensory memory, or echoic memory, is typically characterized by its limited temporal capacity but high informational capacity. Auditory working memory has a more limited informational capacity, and a temporal capacity that can be extended for long periods through active rehearsal. Auditory long-term memory seems to be effectively unlimited in both temporal and informational capacity (Atkinson & Shiffrin, 1968; Kumar et al., 2016; Nees, 2016; Neisser, 1967).

The auditory sequences studied by Barascud et al. used very short cycle lengths, of the order of 1 s, and are therefore likely to fall within the remit of echoic memory. Given that temporal limitations to echoic memory are well-documented in the literature, we might expect these temporal limits to cause the impaired performances observed by Barascud et al. However, some historic work does point to informational limits in echoic memory that can constrain performance in perceptual tasks (Watson, 2016), and such informational limits could also be responsible for Barascud et al.’s observations.

We conducted a behavioral experiment to test these competing explanations. We based this experiment on the regularity detection task from Barascud et al., and created six experimental conditions that orthogonalised two stimulus features: the number of tones in the cycle (10 tones or 20 tones), and the temporal duration of each tone in the cycle (25 ms, 50 ms, or 75 ms). We reasoned that if performance were constrained by informational capacity, then it would be best predicted by the number of tones in the cycle, whereas if performance were constrained by temporal limits, it would be best predicted by the total duration of each cycle. We were particularly interested in the pair of conditions with equal cycle duration but different numbers of tones per cycle (10 × 50 ms = 20 × 25 ms); a decrease in performance in the latter condition would be evidence for informational constraints on regularity detection.

The behavioral results are summarized in Figure 5. Response accuracies are plotted in Figure 5A in terms of the sensitivity metric from signal detection theory. Similar to Barascud et al. (2016), response accuracy was close to ceiling performance across all conditions, with the exception of the condition with the maximum-duration cycles (20 tones each of length 75 ms), where some participants fell away from ceiling performance. Given that accuracies were generally close to ceiling, we instead focus on interpreting reaction-time metrics (Figure 5B). Here we see a clear effect of the number of tones in the cycle, with 10-tone cycles eliciting considerably lower reaction times than 20-tone cycles. This is consistent with the notion of an informational capacity to echoic memory. In particular, comparing the two conditions with equal cycle duration but different numbers of tones per cycle (10 × 50 ms tones; 20 × 25 ms tones), we see that increasing the number of tones substantially impaired performance even when cycle duration stayed constant.

**Figure 5 .**
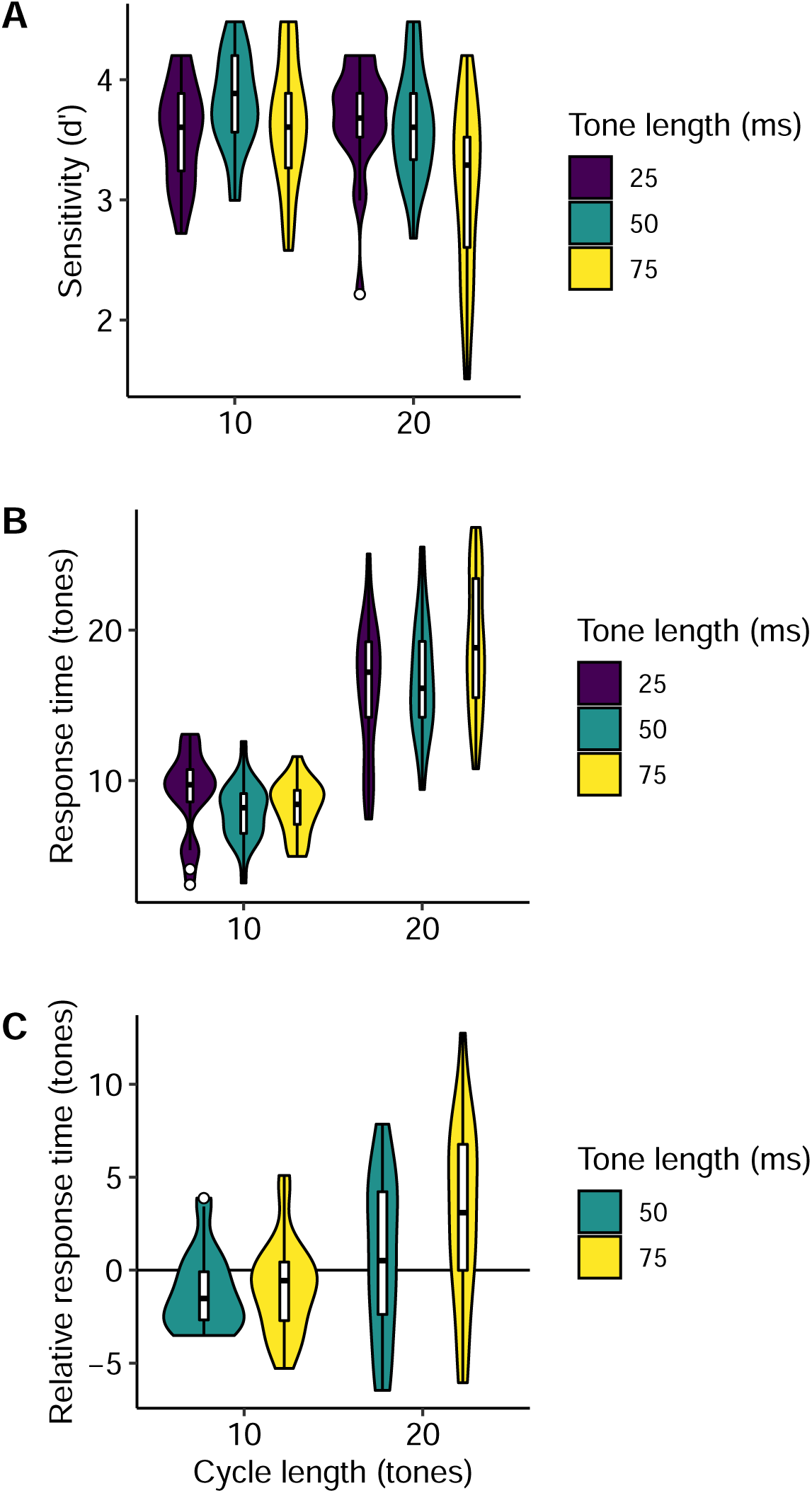
Behavioral results for Experiment 3. **A)** Participant *d*-prime scores by condition, as summarized by violin plots and Tukey box plots. **B)** Participant mean response times by condition, as summarized by violin plots and Tukey box plots. **C)** As **B**, except benchmarking response times against the 25 ms conditions.

Figure 5B does not show a clear effect of tone duration. However, the figure does not account for the repeated-measures structure of the data, meaning that between-condition effects may be partly masked by individual differences between participants. To achieve a more sensitive analysis, Figure 5C takes advantage of the repeated-measures structure of the data, and plots each participant’s response time in the 50-ms and 75-ms conditions relative to their response time in the relevant 25-ms condition. Here we again see null or limited effects of tone duration, except in the case of the maximum-duration condition (20 tones each of length 75 ms), where reaction times seem higher than in the corresponding 25-ms and 50-ms conditions. We tested the reliability of this effect by computing each participant’s difference in mean response time between the 25-ms and 75-ms conditions for the 20-tone cycles, and subtracting the analogous difference in response times for the 10-tone cycles, in other words:

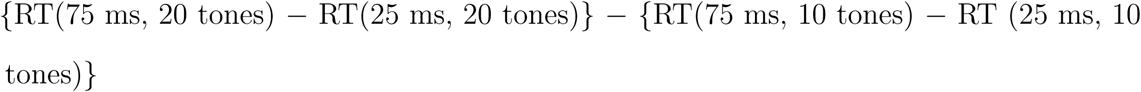

This number summarizes the extent to which increasing tone duration has a stronger effect on reaction times for cycles containing more tones. Using the bias-corrected and accelerated bootstrap (DiCiccio & Efron, 1996), the 95% confidence interval for this parameter was found to be [2.08, 5.93]. The lack of overlap with zero indicates that the effect was fairly reliable: increasing tone duration from 25-ms and 75-ms had a stronger negative effect on reaction times for 20-tone cycles than for 10-tone cycles.

To summarize, then: the behavioral data indicate that performance in this regularity-detection task was primarily constrained by the number of tones in the repeating cycles, rather than their duration. However, the data do suggest a subtle negative effect of tone duration which may manifest for cycles containing large numbers of tones.

We now consider how these effects may be reproduced by incorporating memory effects into the PPM model. Instead of the decay kernel solely operating as a function of time, as in Experiments 1 and 2, it must now account for the number of tones that have been observed by the listener. Various such decay kernels are possible. Here we decided to base our decay kernel on the following psychological ideas, inspired by previous research into echoic memory (Atkinson & Shiffrin, 1968; Nees, 2016; Watson, 2016):

1. Echoic memory operates as a continuously updating buffer that stores recent auditory information.
2. While a memory remains in the buffer, it is represented with high fidelity, and is therefore a reliable source of information for regularity detection mechanisms.
3. The buffer has a limited temporal and informational capacity. Memories will remain in the buffer either until a certain time period has elapsed, or until a certain number of subsequent events has been observed.
4. Once a memory leaves the buffer, it is represented in a secondary memory store.
5. Observations in this secondary memory store contribute less strongly to auditory pattern detection, and gradually decay in salience over time, as in Experiments 1 and 2.

These principles, formalized computationally and applied to the continuous tone sequences from the behavioral experiment, result in the decay kernels described in Figure 6. In each case the buffer is limited to a capacity of 15 tones, which corresponds to a time duration of 0.375 s for 25-ms tones, 0.75 s for 50-ms tones, and 1.125 s for 75-ms tones. While the *n*-gram observation remains within this buffer, its weight is *w*_0_ = 1.0; once the memory exits the buffer its weight drops to *w*_1_ = 0.6, and thereafter decays exponentially to *w*_∞_ = 0 with a half life of *t*_0.5_ = 3.5 s. The precise parameters of this decay kernel come from manual optimization to the behavioral data.

**Figure 6 .**
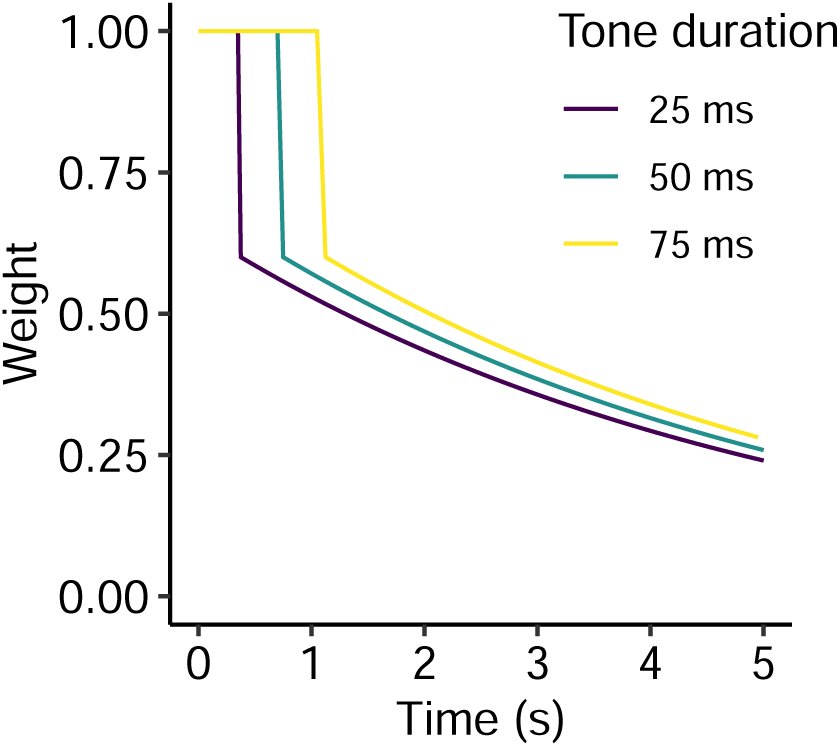
Decay kernels employed in Experiment 3. The temporal duration of the buffer corresponds to the buffer’s informational capacity (15 tones) multiplied by the tone duration.

Weight decay by itself is not sufficient to cause memory loss, because PPM computes its predictions using ratios of event counts, which are preserved under multiplicative weight decay. We therefore introduce stochastic noise to the memory retrieval component of the PPM model, meaning that weight decay reduces the signal-to-noise ratio, and thereby gradually eliminates the memory trace of the original observation. In our optimized model this noise corresponds to a Gaussian with standard deviation *σ*_*ϵ*_ = 0.8.

Applied to an individual trial, the model returns the information content for each tone in the sequence, corresponding to the surprisingness of that tone in the context of the prior portion of the sequence (Figure 7). Following Barascud et al. (2016), we suppose that the listener identifies the transition from random to regular tone patterns by detecting the ensuing drop in information content. We model this process using a non-parametric change-detection algorithm that sequentially applies the Mann-Whitney test to identify changes in a time series’ location while controlling the false positive rate to 1 in 10,000 observations (Ross, Tasoulis, & Adams, 2011).

All stimuli were statistically independent from one another, and so responses should not be materially affected by experiences on previous trials. For simplicity and computational efficiency, we therefore left the PPM-Decay model’s long-term learning weight (*w*_∞_) fixed at zero, and reset the model’s memory store between each trial.

We analyzed 6 different PPM-Decay configurations, aiming to understand how the model’s different features contribute to task performance, and which are unnecessary for explaining the perceptual data. Specifically, we built the proposed model step-by-step from the original PPM model, first adding exponential decay, then adding retrieval noise, then adding the memory buffer. We tested three versions of the final model with different buffer capacities: 5 items, 10 items, and 15 items. We manually optimized each model configuration to align mean participant response times to mean model response times, producing the parameter sets listed in Table 1.

**Table 1.**
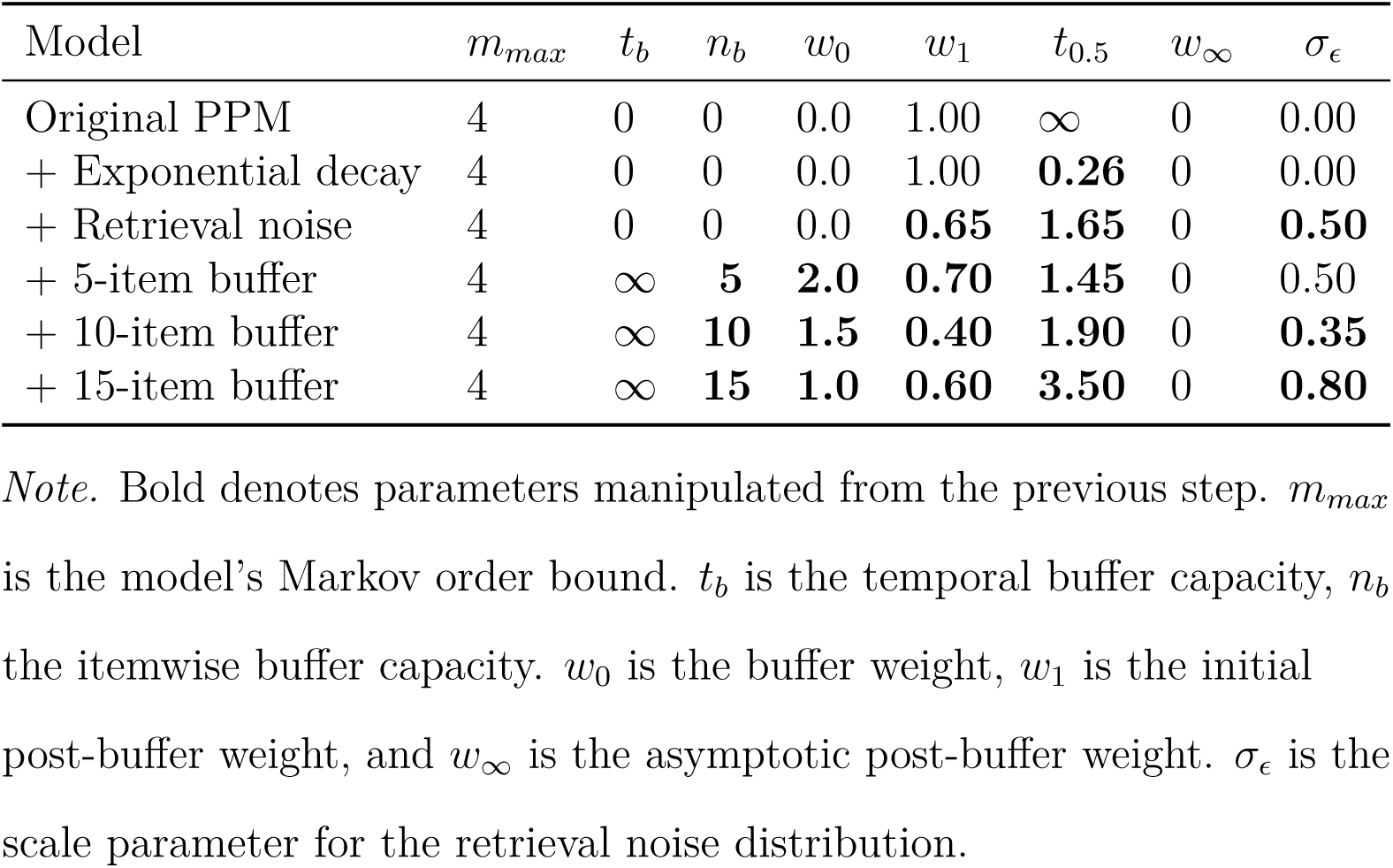
Optimized model parameters for Experiment 3.

| Model | $m_{max}$ | $t_b$ | $n_b$ | $w_0$ | $w_1$ | $t_{0.5}$ | $w_\infty$ | $\sigma_\epsilon$ |
| --- | --- | --- | --- | --- | --- | --- | --- | --- |
| Original PPM | 4 | 0 | 0 | 0.0 | 1.00 | ∞ | 0 | 0.00 |
| + Exponential decay | 4 | 0 | 0 | 0.0 | 1.00 | **0.26** | 0 | 0.00 |
| + Retrieval noise | 4 | 0 | 0 | 0.0 | **0.65** | **1.65** | 0 | **0.50** |
| + 5-item buffer | 4 | ∞ | **5** | **2.0** | **0.70** | **1.45** | 0 | 0.50 |
| + 10-item buffer | 4 | ∞ | **10** | **1.5** | **0.40** | **1.90** | 0 | **0.35** |
| + 15-item buffer | 4 | ∞ | **15** | **1.0** | **0.60** | **3.50** | 0 | **0.80** |

*Note.* Bold denotes parameters manipulated from the previous step. $m_{max}$ is the model’s Markov order bound. $t_b$ is the temporal buffer capacity, $n_b$ the itemwise buffer capacity. $w_0$ is the buffer weight, $w_1$ is the initial post-buffer weight, and $w_\infty$ is the asymptotic post-buffer weight. $\sigma_\epsilon$ is the scale parameter for the retrieval noise distribution.

### Original PPM

As expected, the original PPM model proved not to be sensitive to tone length or to alphabet size (Figure 8A). Furthermore, the model systematically outperformed the participants, with an average reaction time of 6.23 tones compared to the participants’ mean reaction time of 12.90.

**Figure 8 .**
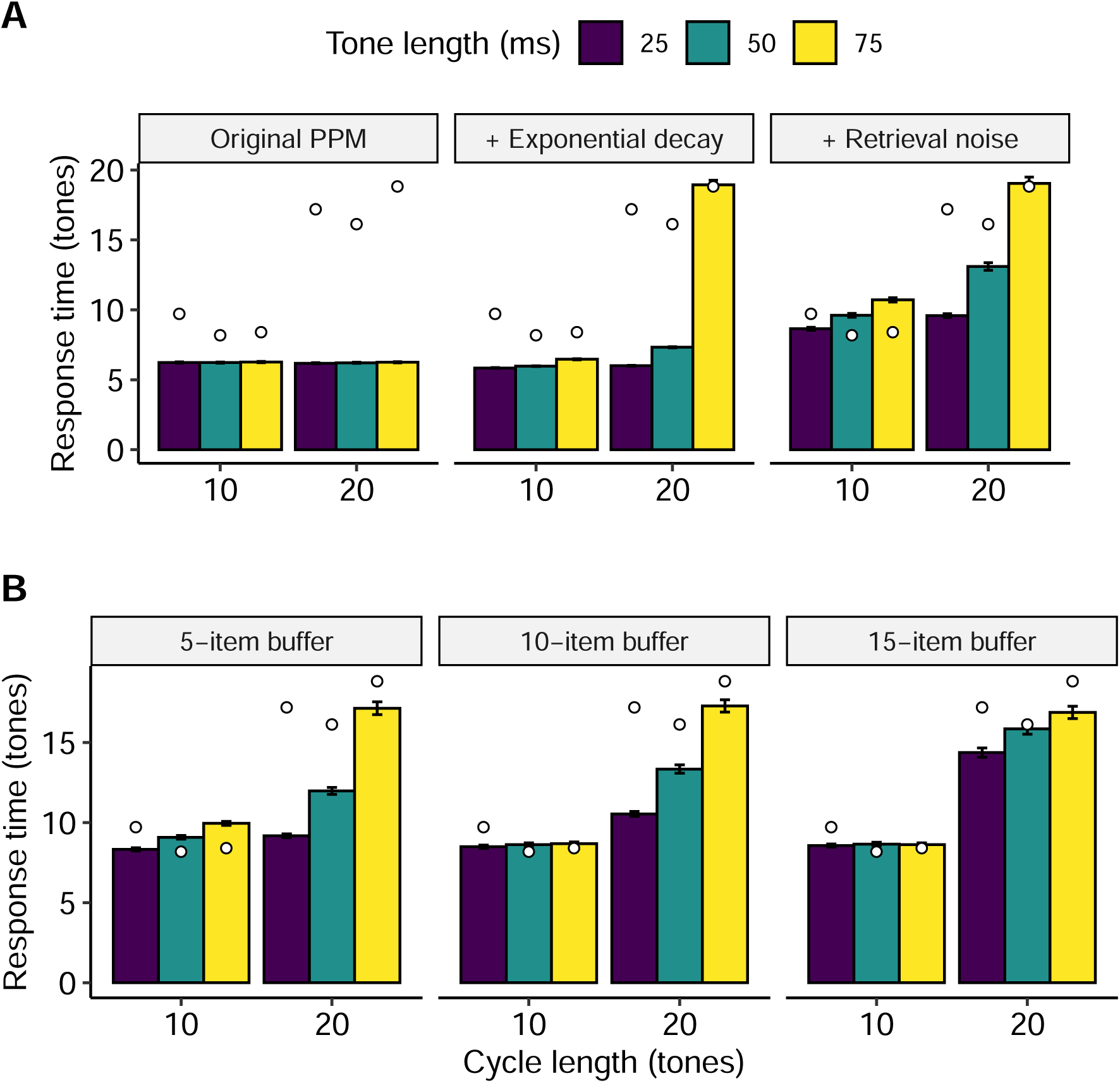
Modeling participant data (mean response times, white circles) with different model configurations (mean simulated response times, solid bars). Error bars denote 95% confidence intervals computed using the central limit theorem. **A)** Progressively adding exponential weight decay and retrieval noise to the original PPM model. **B)** Progressively adding longer buffers to the PPM-Decay model.

### Adding exponential decay

Here we add time-based exponential decay, as in Experiments 1 and 2. One might expect this feature to induce a negative relationship between pattern-detection performance and cycle length. We do observe such an effect, but only with a very fast memory-decay rate (half life = 0.26 s; Figure 8A). This robustness of models without retrieval noise to memory decay can be rationalized by observing that, even as absolute weights of memory traces decrease with memory decay, the important information, namely the ratios of these weights, remains more or less preserved, and so the pattern-detection algorithm continues to perform well. Further to this, the model is problematic in that it substantially outperforms participants in the 10-tone conditions, and exhibits no clear discontinuity in performance between the 10-tone conditions and the 20-tone conditions.

### Adding retrieval noise

Retrieval noise increases the model’s sensitivity to memory decay, and means that the drop in performance from the shortest cycles (10 tones, 25 ms/tone) to the longest cycles (20 tones, 75 ms/tone) can be replicated with a more plausible half-life of 1.65 s (Figure 8A). However, the model still fails to capture the discontinuity in reaction times between 10-tone and 20-tone conditions, especially with tone lengths of 25 and 50 ms.

### Adding the memory buffer

We anticipated that a buffer with an informational capacity limit between 10 tones and 20 tones should be able to replicate the behavioral discontinuity between 10-tone and 20-tone conditions. The 10-tone cycles should largely fit in such a buffer, resulting in near-ceiling performance in the 10-tone conditions; conversely, the 20-tone cycles should be too big for the buffer, resulting in performance deterioration. Figure 8B shows that such an effect does indeed take place with a 15-tone buffer. In contrast, shorter buffers (5 tones, 10 tones) do not elicit this clear discontinuity between 10-tone and 20-tone conditions. The resulting model also replicates the insensitivity to tone duration in the 10-tone conditions, and the adverse effect of increasing tone duration to 75 ms in the 20-tone condition that was hinted at in the behavioral data. It therefore seems clear that a PPM-Decay model with a finite-capacity buffer can explain the main patterns of reaction times observed in this experiment, in contrast to the original PPM model.

## Discussion

PPM is a powerful sequence prediction algorithm that has proved well-suited to modeling the cognitive processing of auditory sequences (Agres et al., 2018; Barascud et al., 2016; Egermann et al., 2013; Pearce & Müllensiefen, 2017; Pearce et al., 2010; Pearce & Wiggins, 2006). In these contexts, PPM has traditionally been interpreted as an ideal observer, simulating an (approximately) optimal strategy for predicting upcoming auditory events on the basis of learned statistics. This modeling strategy has proved very useful for elucidating the role of statistical cognition in auditory perception (Barascud et al., 2016; Pearce, 2018).

Here we introduced a customizable decay kernel to PPM, which downweights historic observations as time passes and subsequent observations are registered in memory. This decay kernel is useful for two primary reasons. First, it makes PPM a better approximation to an ideal observer when the underlying sequence statistics change over time, as is common in many real-world listening contexts. Second, it allows the model to capture the multi-stage nature of human auditory memory, with its corresponding capacity limitations and temporal profiles.

We applied this new PPM-Decay model in three experiments. The first experiment analyzed sequences generated from a statistical model whose underlying parameters evolved over time, and verified that PPM-Decay better approximates an ideal observer than PPM when applied to such sequences. The second experiment simulated a musically naive listener who gradually learns to predict chord progressions through exposure to compositions from three musical styles: popular music, jazz music, and chorale harmonizations by J. S. Bach. Again, we found that PPM-Decay better approximated an ideal observer than the original PPM model. The ideal model configuration incorporated a recency effect, reflecting how the underlying statistics of the chord progressions differ between compositions, and evolve during the course of individual compositions. However, the model’s decay kernel also incorporated a positive asymptote, allowing the model to develop long-term knowledge of certain statistical regularities that are shared between different compositions from the same musical style.

The third experiment revisited an auditory detection paradigm from Barascud et al. (2016), where participants had to detect transitions between random and regular sections in tone sequences that varied in alphabet size and tone length. Barascud et al. found tentative evidence for auditory pattern detection being constrained by the capacity limitations of echoic memory, but were unable to determine whether these results reflected temporal limitations (e.g. echoic memory only spans two seconds) or informational limitations (e.g. echoic memory can only hold up to 15 tones). We conducted a new behavioral experiment using stimuli designed to distinguish these two possibilities, by varying tone duration and the number of tones in the regular patterns independently. The resulting data implied that human performance stayed constant as long as the relevant auditory input could fit within a buffer of limited itemwise capacity. We formalized this explanation computationally with our PPM-Decay model, and showed that the model could successfully reproduce the observed behavioral data, in contrast to simpler model variants such as the original PPM model (Barascud et al., 2016; Bunton, 1997; Pearce, 2005) or a PPM model with solely exponential memory decay.

We anticipate that this PPM-Decay model should prove useful for other applications in auditory modeling. The combination of the statistical power of PPM and the flexible decay kernel makes the model well-suited to simulating online auditory statistical learning under memory constraints and in changing statistical environments. A particularly relevant application domain is music cognition, which has already made significant use of PPM models without decay kernels (Di Giorgi, Dixon, Zanoni, & Sarti, 2017; Egermann et al., 2013; Harrison & Pearce, 2018; Pearce & Müllensiefen, 2017; Pearce et al., 2010; Pearce & Wiggins, 2006). Incorporating decay kernels into these models should be useful for capturing how recency effects and memory limitations influence the probabilistic processing of musical structure. However, the PPM-Decay algorithm itself is relatively domain-agnostic, and should be applicable to any sequential domain where observations can be approximated as discrete symbols drawn from a finite alphabet. We anticipate that our publicly available R package “ppm” should prove useful for supporting such work (https://github.com/pmcharrison/ppm).

An important avenue for future work is to improve our understanding of the ideal decay kernels for different modeling applications. When optimizing a decay kernel for predictive performance on a corpus of sequences, we learn about the statistical structure of that corpus, specifically the sense in which historical events of different vintages contribute useful information about upcoming events. Such analyses are particularly relevant to computational musicology, where a common goal is to quantify statistical processes underlying music composition. When optimizing a decay kernel to reproduce human performance, we learn about the predictive strategies actually used by humans, and the sense in which they may be constrained by cognitive limitations. The optimized decay kernel from Experiment 3 provides an initial model that seems to account well for the behavioral data collected here, but further empirical work is required to constrain the details of this model and to establish its generalizability to different experimental contexts.

A primary limitation of the PPM and PPM-Decay models is that they operate over discrete representations, and do not model the process by which these discrete representations are extracted from the auditory signal. This simplification is convenient when modeling systems such as music and language, which are often well-suited to symbolic expression, but it is problematic when modeling continuous stimulus spaces. One solution to this problem is to adopt continuous-input models (e.g. Skerritt-Davis & Elhilali, 2018, 2019), where discretization plays no part; however, such models typically struggle to capture the kinds of structural dependencies common in music and language, and do not reflect the apparent importance of categorical perception in human auditory perception (e.g. Repp, 1984). One alternative way forward might be to prefix the PPM-Decay model with an unsupervized discretization algorithm, such as *k*-means clustering (Steinley, 2006).

The PPM-Decay algorithm can become computationally expensive with long input sequences. In the naive implementation, the algorithm must store an explicit record of each *n*-gram observation as it occurs, meaning that the time and space complexity for generating a predictive distribution is linear in the length of the training sequence. However, particular families of decay kernels can support more efficient implementations. For example, a decay kernel comprising the sum of *N* exponential functions can be implemented as a set of *N* counters for each *n*-gram, each of which is incremented upon observing the respective *n*-gram, and each of which is decremented by a fixed ratio at each timestep. This implementation has bounded time and space complexity as regards the length of the training sequence. Such approaches should be useful for speeding the application of the PPM-Decay model to large datasets, and for improving its biological plausibility.

The PPM and PPM-Decay models assume that listeners process auditory stimuli by computing transition probabilities from memories of *n*-gram observations. While *n*-gram models seem to provide a good account of auditory processing (Barascud et al., 2016; Pearce, 2018), they may not be sufficient to explain all aspects of auditory learning. For example, *n*-gram models struggle to explain how listeners can (albeit with some difficulty) learn non-adjacent dependencies (Endress, 2010; Wilson et al., 2018) or recursive grammatical structures (Rohrmeier & Cross, 2009; Rohrmeier et al., 2012). Some of these phenomena might be explained by incorporating further modifications to the memory model; for example, non-adjacent dependencies could be learned by combining *n*-gram modeling with the abstraction method of Thiessen & Pavlik (2013). Other phenomena, such as the acquisition of recursive grammars, might only be explained by alternative modeling approaches. This remains a challenge for future research.

Several alternative cognitive models of sequence prediction have explicitly Bayesian formulations (e.g. Skerritt-Davis & Elhilali, 2018; Bröker et al., 2018; Meyniel et al., 2016). This approach is appealing because it formally motivates the predictive algorithm from a set of assumptions about the underlying sequence statistics. Such approaches can also be applied to mixed-order Markov models such as PPM, but typically they come with substantially increased computational complexity (Teh, 2006), which may prove impractical for many cognitive modeling applications. Nonetheless, it would be worth examining how the present approaches might be motivated as computationally efficient approximations to Bayes-optimal models.

## Methods

### Model

Our PPM-Decay model embodies a predictive processing account of auditory regularity detection. It supposes that listeners acquire an internal model of incoming sounds through automatic processes of statistical learning, and use this model to generate prospective predictions for upcoming auditory events. The model derives from the PPM algorithm (Bunton, 1997; Cleary & Witten, 1984), but adds three psychological principles:

a. The memory salience of a given observation decays as a function of the timepoints of subsequently observed events and the timepoint of memory retrieval.
b. There exists some noise, or uncertainty, in memory retrieval.
c. A limited-capacity memory buffer constrains learning and prediction. Contiguous events (*n*-grams) must fit into this buffer to be internalized or to contribute to prediction generation.

Each of these three features can be enabled or disabled in isolation. In ideal-observer analyses, such as Experiments 1 and 2, it often makes sense to omit features b) and c), becaues they correspond to cognitive constraints that typically impair prediction. Here we therefore omit these two features for the ideal-observer analyses (Experiments 1 and 2), but retain them for the behavioral analyses in Experiment 3.

Many variants of PPM exist in the literature (Bunton, 1997; Cleary & Teahan, 1997; Cleary & Witten, 1984; Moffat, 1990). Our formulation incorporates the interpolated smoothing technique of Bunton (1997), but avoids techniques such as exclusion, update exclusion, and state selection, because they do not generalize naturally to decay-based models.

#### Domain

The model assumes that the auditory input can be represented as a sequence of symbols drawn from a discrete alphabet; the cognitive processes involved in developing this discrete representation are not addressed here. Let 𝒜 denote the discrete alphabet, let () denote an empty sequence, and let 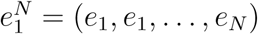 denote a sequence of *N* symbols, where *e*_*i*_ ∈ 𝒜 is the *i*th symbol in the sequence, and 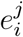 is defined as

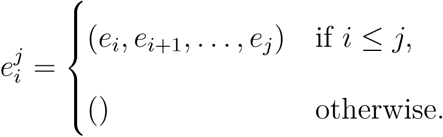

We suppose that this sequence is presented over time, and denote the timepoint of the *i*th symbol as *τ*_*i*_.

Now suppose that 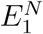 is a random variable corresponding to a sequence of length *N*. We consider an observer predicting each symbol of 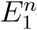 based on the previously observed symbols. This corresponds to the probability distribution 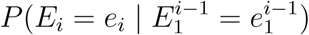, which we will abbreviate as 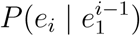. The model is tasked with estimating this conditional probability distribution.

#### Learning

The model learns by counting occurrences of different sequences of length *n* termed *n-grams* (*n* ∈ ℕ^+^), where *n* is termed the *n-gram order.* As in PPM, the model counts *n*-grams for all *n* ≤ *n*_*max*_ (*n*_*max*_ ∈ ℕ^+^), where *n*_*max*_ is the *n-gram order bound*. A three-symbol sequence (*e*_1_, *e*_2_, *e*_3_) contains six *n*-grams: (*e*_1_), (*e*_2_), (*e*_3_), (*e*_1_, *e*_2_), (*e*_2_, *e*_3_), and (*e*_1_, *e*_2_, *e*_3_).

We suppose that *n*-grams are extracted from a finite-capacity buffer (Figure 9). Successive symbols enter and leave this buffer in a first-in first-out arrangement, so that the buffer represents a sliding window over the input sequence. The buffer has two capacity limitations: *itemwise capacity* and *temporal capacity*. The itemwise capacity, *n*_*b*_, determines the maximum number of symbols stored by the buffer; the temporal capacity, *t*_*b*_, determines the maximum amount of time that a given symbol can remain in the buffer before expiry. Generally speaking, itemwise capacity will be the limiting factor at fast presentation rates, whereas temporal capacity will be the limiting factor at slow presentation rates. As *n*-grams may only be extracted if they fit completely within the buffer, these capacities bound the order of extracted *n*-grams. Correspondingly, we constrain *n*_*max*_ (the *n*-gram order bound) not to exceed *n*_*b*_ (the itemwise buffer capacity).

**Figure 9 .**
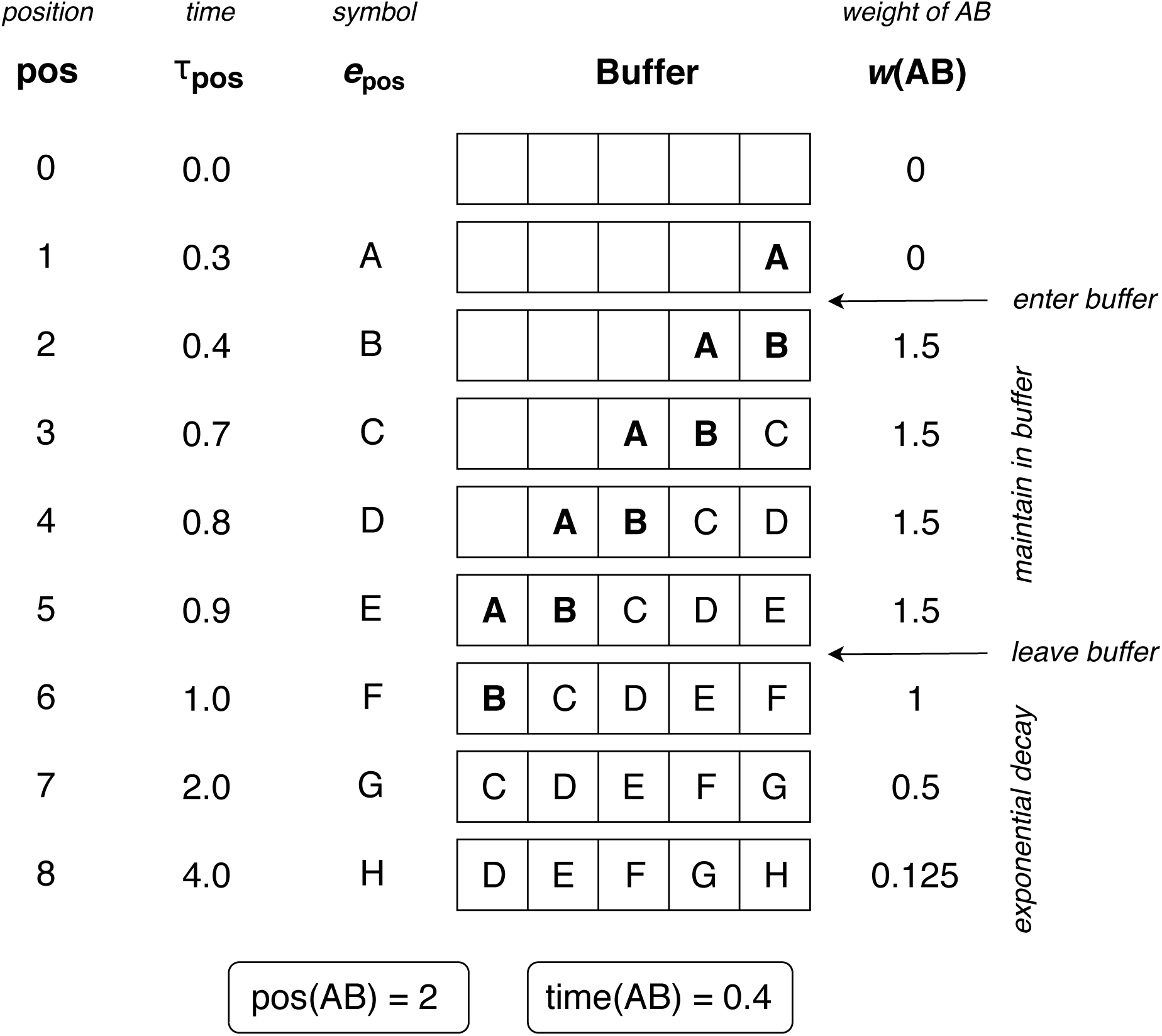
Schematic figure illustrating the accumulation of observations within a memory buffer with an itemwise capacity of 5. Weights for the *n*-gram “AB” are displayed as a function of time, assuming a buffer weight (*w*_0_) of 1.5, an initial post-buffer weight (*w*_1_) of 1, a half life (*t*_0.5_) of 1 second, and an asymptotic post-buffer weight (*w*_∞_) of 0.

In PPM, *n*-gram observations are recorded by incrementing a counter. Our PPM-Decay model also stores the ordinal position within the input sequence when the observation occurred; this is necessary for simulating the temporal dynamics of auditory memory. For each *n*-gram *x*, we define count(*x*) as the total number of observations of *x*, and pos(*x*) as a list of ordinal positions in the input sequence when these observations occurred, defined with respect to the final symbol in the *n*-gram. pos(*x*) is initialized as an empty list; each time a new *n*-gram *x* is observed, the respective ordinal position is appended to the list. count(*x*) is then represented implicitly as the length of pos(*x*).

The input sequence is processed one symbol at a time, from beginning to end. Observing the *i*th symbol, *e*_*i*_, yields up to *n*_*max*_ *n*-gram observations, corresponding to all *n*-grams in the buffer that terminate with the most recent symbol: 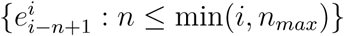. If the buffer component of the model is enabled, an *n*-gram observation will only be recorded if it fits completely within the itemwise and temporal capacities of the buffer; the former constraint is ensured by the constraint that *n*_*max*_ ≤ *n*_*b*_, but the latter must be checked by comparing the current timepoint (corresponding to the final symbol in the *n*-gram) with the timepoint of the first symbol of the *n*-gram. If the current ordinal position is written pos_end_, and the *n*-gram length is written size(*x*), then the necessary and sufficient condition for *n*-gram storage is

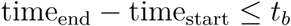

where

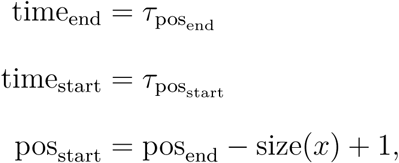

*τ*_*i*_ is the *i*th timepoint in the input sequence, and *t*_*b*_ is the temporal buffer capacity, as before. Table 2 describes the information potentially learned from training on the sequence (*a, b, a*).

**Table 2.**
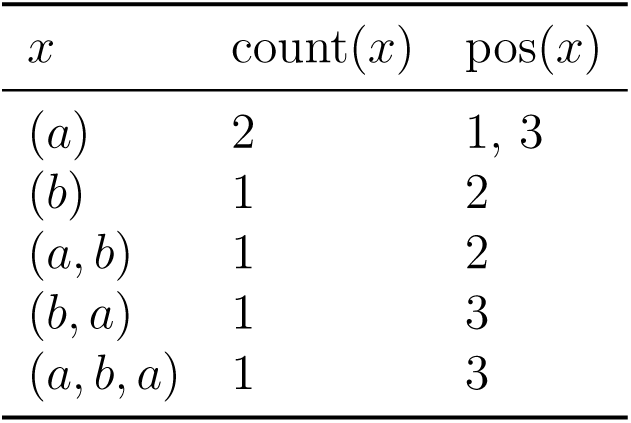
n*-grams learned from training on the sequence* (*a, b, a*).

| $x$ | count($x$) | pos($x$) |
| --- | --- | --- |
| $(a)$ | 2 | 1, 3 |
| $(b)$ | 1 | 2 |
| $(a, b)$ | 1 | 2 |
| $(b, a)$ | 1 | 3 |
| $(a, b, a)$ | 1 | 3 |

#### Memory decay

In the original PPM algorithm, the influence of a given *n*-gram observation is not affected by the passage of time or the encoding of subsequent observations. This contrasts with the way in which human observers preferentially weight recent observations over historic observations (Bröker et al., 2018; Harrison, 2011; Mattar et al., 2016; Meyniel et al., 2016; O’Reilly, 2013; Squires et al., 1976; Yu & Cohen, 2008). This inability to capture recency effects limits the validity of PPM as a cognitive model.

Here we address this problem. We suppose that the influence, or *weight*, of a given *n-*gram observation varies as a function both of the current timepoint and the timepoints of the symbols that have since been observed. This weight decay function represents the following hypotheses about auditory memory:

a. Each *n*-gram observation begins in the memory buffer (Figure 9). Within this buffer, observations do not experience weight decay.
b. Upon leaving the buffer, observations enter a secondary memory store. This transition is accompanied by an immediate drop in weight.
c. While in the secondary memory store, observations experience continuous weight decay over time, potentially to a non-zero asymptote.

These hypotheses must be considered tentative, given the scarcity of empirical evidence directly relating memory constraints to auditory prediction. However, the notion of a short-lived memory buffer is consistent with pre-existing concepts of auditory sensory memory (Atkinson & Shiffrin, 1968; Nees, 2016; Neisser, 1967), and the continuous-decay phenomenon is consistent with well-established recency effects in statistical learning (Bröker et al., 2018; Harrison, 2011; Mattar et al., 2016; Meyniel et al., 2016; O’Reilly, 2013; Squires et al., 1976; Yu & Cohen, 2008).

We formalize these ideas as follows. For readability, we write pos(*x, i*) for the *i*th element of pos(*x*), corresponding to the ordinal position of the *i*th observation of *n*-gram within the input sequence, defined with respect to the final symbol of the *n*-gram. Similarly, we write time(*x, i*) as an abbreviation of *τ*_pos(*x,i*)_, the timepoint of the *i*th observation of *n*-gram *x*. We then define *w*(*x, i, t*) as the weight for the *i*th observation of *n*-gram *x* for an observer situated at time *t*:

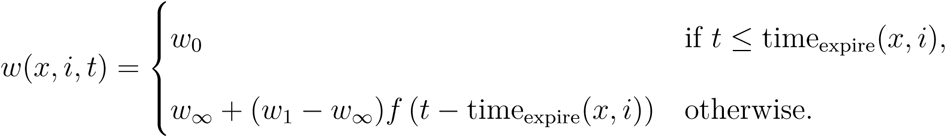

Here *w*_0_ is the *buffer weight, w*_1_ is the *initial post-buffer weight*, and *w*_∞_ is the *asymptotic post-buffer weight* (*w*_0_ ≥ *w*_1_ ≥ *w*_∞_ ≥ 0). The function *f* defines an exponential decay with half-life equal to *t*_0.5_, with *t*_0.5_ > 0:

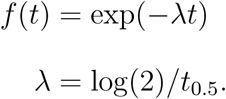

time_expire_(*x, i*) denotes the timepoint at which the *i*th observation of *n*-gram *x* expires from the buffer, computed as the earliest point when either the temporal capacity or the itemwise capacity expires. The temporal capacity expires when *t*_*b*_ seconds have elapsed since the first symbol in the *n*-gram, whereas the itemwise capacity expires when *n*_*b*_ symbols have been observed since the first symbol in the *n*-gram:

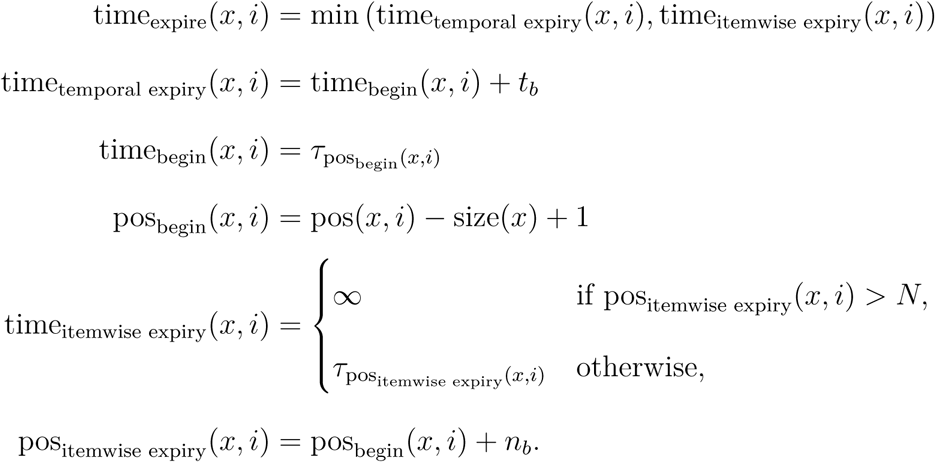

An illustrative memory-decay profile is shown in Figure 10.

**Figure 10 .**
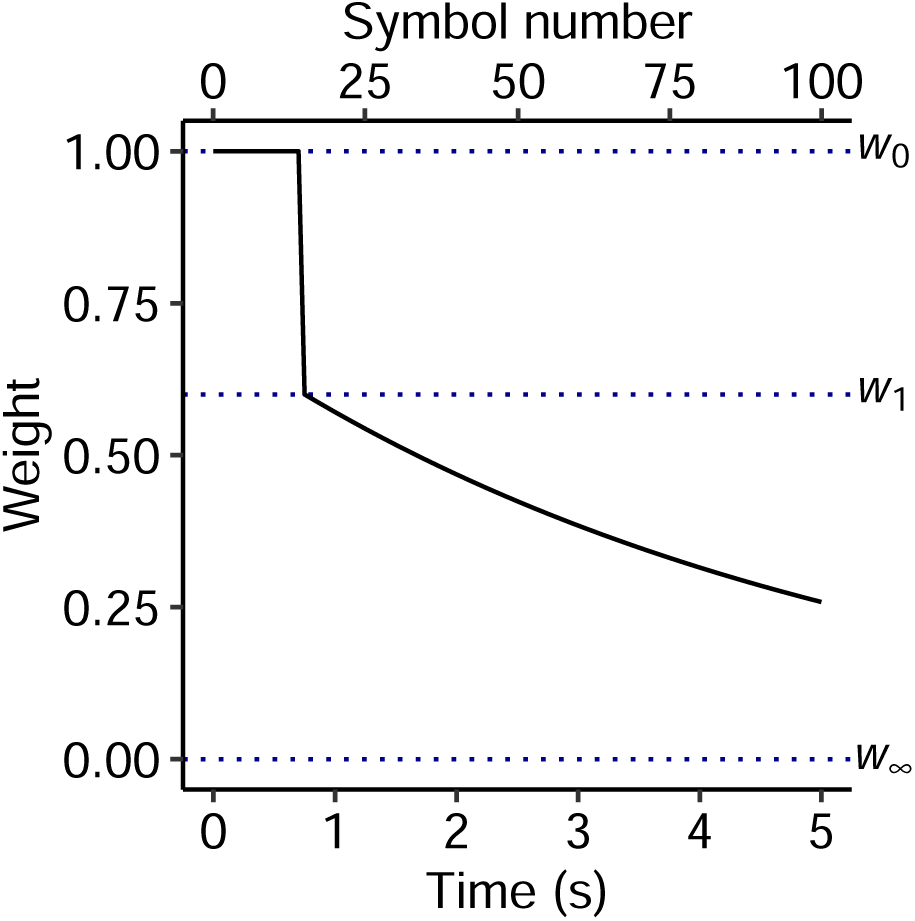
Weight decay for an *n*-gram of length one plotted as a function of relative observer position, assuming that new symbols continue to be presented every 0.05 seconds. Model parameters are set to *t*_*b*_ = 2, *n*_*b*_ = 15, *w*_0_ = 1.0, *t*_0.5_ = 3.5, *w*_1_ = 0.6, and *w*_∞_ = 0, as optimized in Experiment 3.

Memory traces accumulate over repeated observations of the same *n*-gram. We define *W*(*x, t*), the accumulated weight for an *n*-gram *x*, as

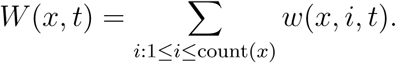

As currently specified, memory decay does not necessarily cause forgetting, because the same information may be preserved in the ratios of *n*-gram weights even as the absolute values of the weights shrink. For example, consider a pair of *n*-grams *AB* and *AC* with weights 4 and 1 respectively, both undergoing exponential decay to an asymptotic weight of 0. From these *n*-gram weights, the model can estimate the probability that *B* follows *A* as *p*(*B* | *A*) = 4*/*(4 + 1) = 0.8. After one half-life, the new counts are 2 and 0.5 respectively, but the maximum-likelihood estimate remains unchanged: *p*(*B* | *A*) = 2*/*(2 + 0.5) = 0.8.

A better account of forgetting can be achieved by supposing that memory traces must compete with noise factors introduced by imperfections in auditory memory; in this case, shrinking the absolute values of *n*-gram weights decreases their signal-to-noise ratio and hence induces forgetting. Here we model imperfections in memory retrieval by adding truncated Gaussian noise to the retrieved weights:

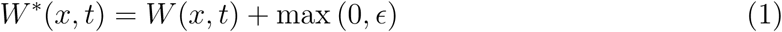

where *W*\*(*x, t*) is the retrieved weight of *n*-gram *x* at time *t*, and 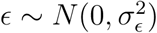 represents Gaussian noise uncorrelated across *n*-grams or timepoints. Setting 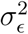 to zero disables the noise component of the model.

#### Prediction

Traditionally, a maximum-likelihood *n*-gram model estimates the probability of symbol *e*_*i*_ given context 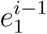 by taking all *n*-grams beginning with 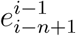 and finding the proportion that continued with *e*_*i*_. For *n* ≤ *i*:

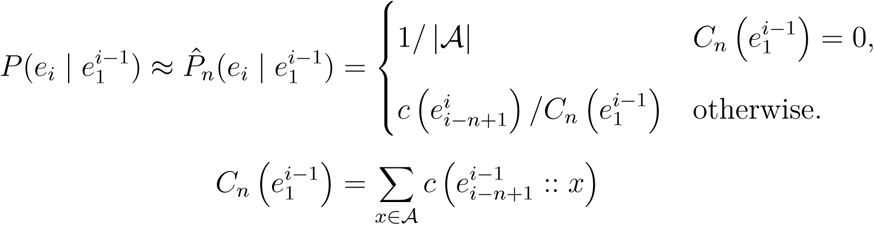

where 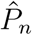 denotes an *n*-gram probability estimator of order 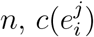 is the number of times *n*-gram 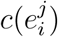 occurred in the training set, and 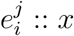 denotes the concatenation of sequence 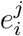 and symbol *x*. The *n*-gram model predicts from the previous *n* − 1 symbols, and therefore constitutes an (*n* − 1)th-order Markov model. Note that the estimator defaults to a uniform distribution if 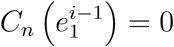, when the context has never been seen before. Note also that the predictive context of a 1-gram model is the empty sequence 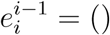.

To incorporate memory decay into a maximum-likelihood *n*-gram model, we replace the count function *c* with the retrieval weight function *W*\*. For *n* ≤ *i*:

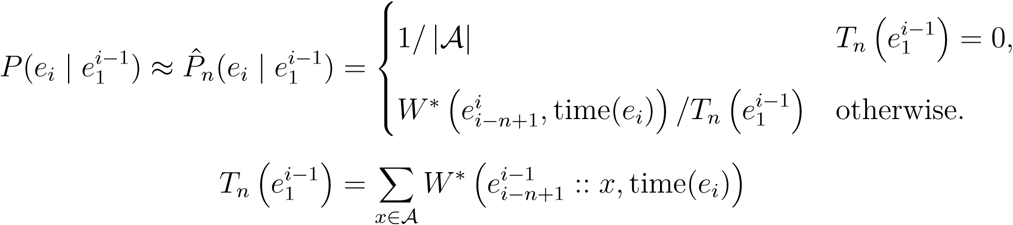

This decay-based model degenerates to the original maximum-likelihood model when *w*_0_ = 1, *t*_*b*_ → ∞, *n*_*b*_ → ∞, *σ*_*ϵ*_ = 0 (i.e. an infinite-length memory buffer with unit weight and no retrieval noise).

High-order *n*-gram models take into account more context when generating their predictions, and are hence capable of greater predictive power; however, this comes at the expense of greater tendency to overfit to training data. Conversely, low-order models are more robust to overfitting, but this comes at the expense of lower structural specificity. Smoothing techniques combine the benefits of both high-order and low-order models by merging *n*-gram models of different orders, with model weights varying according to the amount of training data. Here we use interpolated smoothing as introduced by Bunton (1996, 1997). For *n* ≤ *i*, the unnormalized interpolated *n*-gram estimator is recursively defined as a weighted sum of the *n*th-order maximum-likelihood estimator and the (*n* − 1)th-order interpolated estimator:

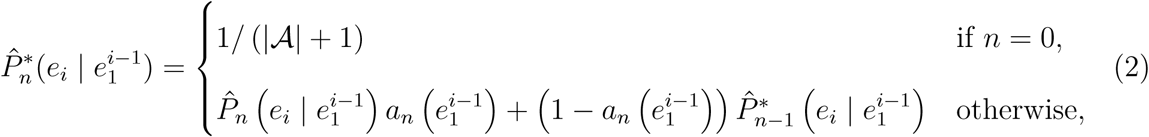

where 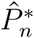 is the *n*th-order unnormalized interpolated *n*-gram estimator, 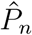 is the *n*th-order maximum-likelihood estimator, |𝒜| is the alphabet size, and *a*_*n*_ is a function of the context sequence that determines how much weight to assign to 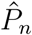, the maximum-likelihood *n*-gram estimator of order *n*.

The unnormalized interpolated estimator defines an improper probability distribution that does not necessarily sum to 1. We therefore define 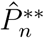 as the normalized interpolated estimator:

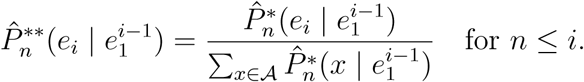

Note that the need for normalization can alternatively be avoided by redefining 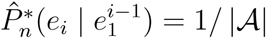 for *n* = 0 in Equation (2), meaning that the interpolated smoothing terminates with a proper probability distribution. However, we keep the original definition to preserve equivalence with Bunton (1997) and Pearce (2005).

The weighting function *a*_*n*_ corresponds to the so-called “escape mechanism” of the original PPM algorithm. Pearce & Wiggins (2004) review five different escape mechanisms, termed “A” (Cleary & Witten, 1984), “B” (Cleary & Witten, 1984), “C” (Moffat, 1990), “D” (Howard, 1993), and “AX” (Moffat, Neal, & Witten, 1998) (see also Bunton, 1996, 1997), each corresponding to different weighting functions *a*_*n*_. Of these, “C” tends to perform the best in data compression benchmarks (Pearce & Wiggins, 2004). However, methods “B”, “C”, “D”, and “AX” do not generalize naturally to decay-based models; in particular, it is difficult to ensure that the influence of an observation is a continuous function of its retrieved weight *w*\*. We therefore adopt mechanism “A”.

In its original formulation, mechanism “A” gives the higher-order model a weight of *a*_*n*_ = 1 − 1*/*(1 + *T*_*n*_), where *T*_*n*_ is the number of times the predictive context has been seen before (which can be interpreted as the observer’s familiarity with the preceding sequence of *n* − 1 tokens). When the context has never been seen before, *T*_*n*_ = 0 and *a*_*n*_ = 0, and the estimator relies fully on the lower-order models; as *T*_*n*_ → ∞, *a*_*n*_ → 1, and the estimator relies fully on the highest-order model. In the original PPM algorithm, the number of times that the predictive context has been seen before is equal to the sum of the weights (or counts) for each possible continuation:

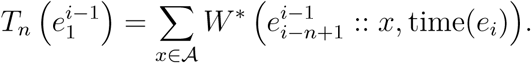

Introducing memory-decay reduces the weights for these prior observations, decreasing the model’s effective experience, and preferentially weighting lower-order models, as might be expected. However, retrieval noise is problematic, because it positively biases the retrieved weights (see Equation (1)), causing the algorithm to overestimate its familiarity with its predictive context, and to overweight high-order predictive contexts as a result. We compensate for this by subtracting the expected value of the retrieval noise’s contribution to *T*_*n*_, which can be computed from standard results for the truncated normal distribution as 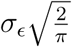, and truncating at zero:

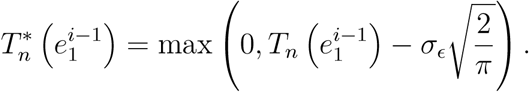

Putting this together, we have (for *i* ≥ *n*):

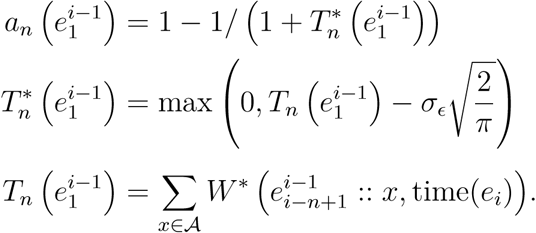

For its final output, the model selects the maximum-order available normalized interpolated estimator. The available orders are constrained by three factors:

a. The *n*-gram order bound: the model cannot predict using *n*-grams larger than *n*_*max*_.
b. The sequence: the predictive context must fit within the observed sequence.
c. The buffer: the predictive context must fit within the buffer at the point when the incoming symbol is observed.

Putting this together, the selected *n*-gram order for generating predictions from a context of 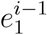 becomes:

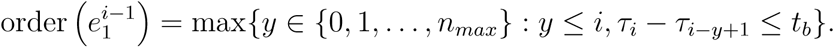

The final model output is then:

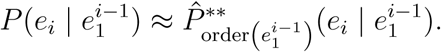

The *n*-gram order bound, *n*_*max*_, constrains the length of *n*-grams that are learned by the model. However, it is often more convenient to speak in terms of the model’s *Markov order, m*_*max*_, defined as the number of preceding symbols that contribute towards prediction generation. A single *n*-gram model generates predictions with a Markov order of *n* − 1; correspondingly, *m*_*max*_ = *n*_*max*_ − 1.

Figure 11 illustrates the interpolated smoothing mechanism. Here we imagine that a model with a Markov order bound of two processes the sequence “abracadabra”, one letter at a time, and then tries to predict the next symbol. The highest-order interpolated distribution, at a Markov order of two, is created by averaging the order-2 maximum-likelihood distribution with the order-1 interpolated distribution, which is itself created by averaging the order-1 maximum-likelihood distribution with the order-0 interpolated distribution. The resulting interpolated distribution combines information from maximum-likelihood models at every order.

**Figure 11 .**
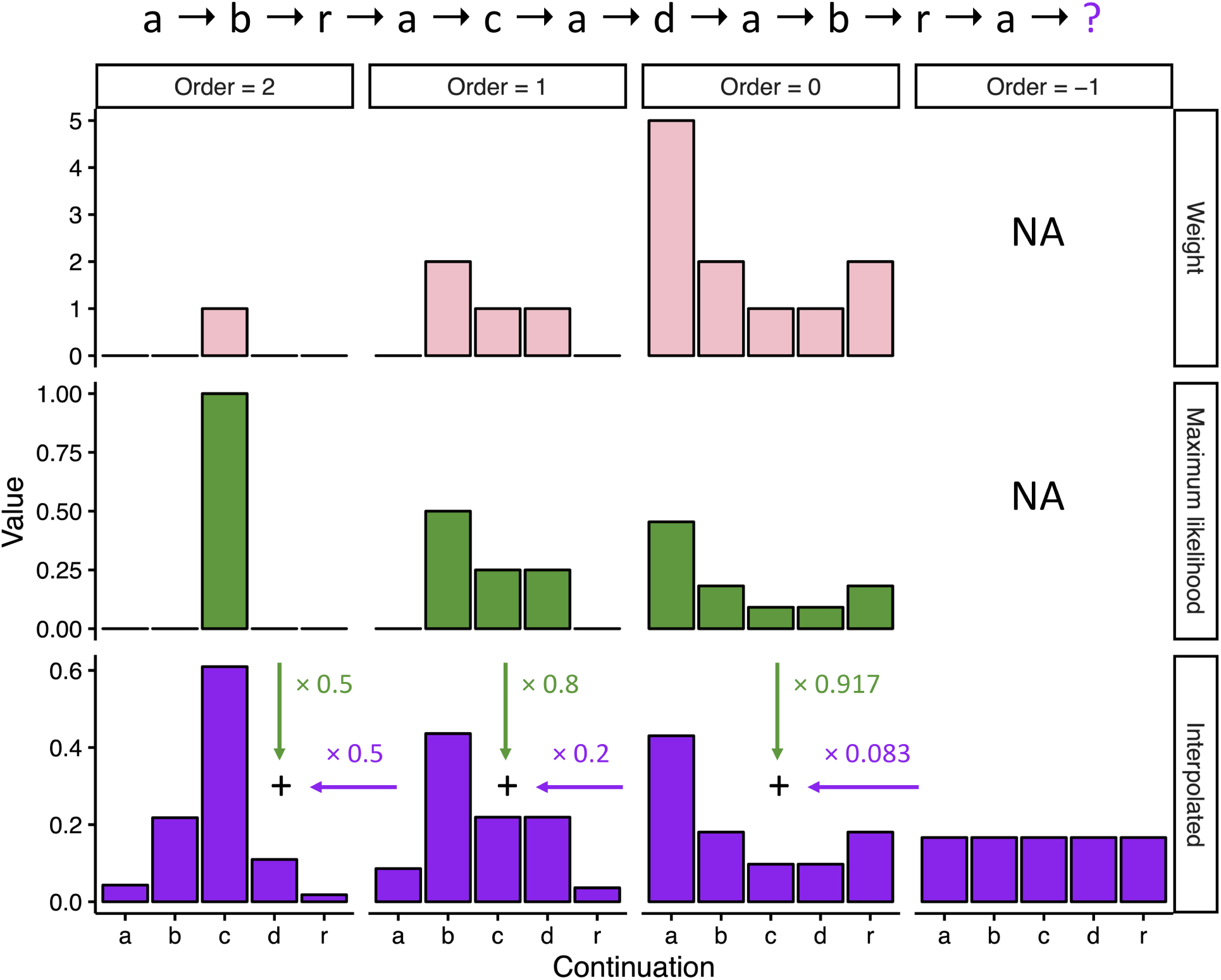
Illustration of the interpolated smoothing mechanism, which blends together maximum-likelihood *n*-gram models of different orders. Here the Markov order bound is two, the predictive context is “abracadabra”, and the task is to predict the next symbol. Columns are identified by Markov order; rows are organized into weight distributions, maximum-likelihood distributions, and interpolated distributions. Maximum-likelihood distributions are created by normalizing the corresponding weight distributions. Interpolated distributions are created by recursively combining the current maximum-likelihood distribution with the next-lowest-order interpolated distribution. The labelled arrows give the weight of each distribution, as computed using escape method “A”. The “Order = −1” column identifies the termination of the interpolated smoothing, and does not literally mean a Markov order of −1.

We have implemented the resulting model in a freely available R package, “ppm”, the core of which is written in C++ for speed. With this package, it is possible to define a PPM-Decay model customized by the eight hyperparameters summarized in Table 3. The package also supports simpler versions of PPM-Decay, where (for example) the buffer functionality is disabled but the exponential-decay functionality is preserved. The resulting models can then be evaluated on arbitrary symbolic sequences. The package may be accessed from its open-source repository at https://github.com/pmcharrison/ppm or its permananent archive at https://doi.org/10.5281/zenodo.2620414.

**Table 3.**
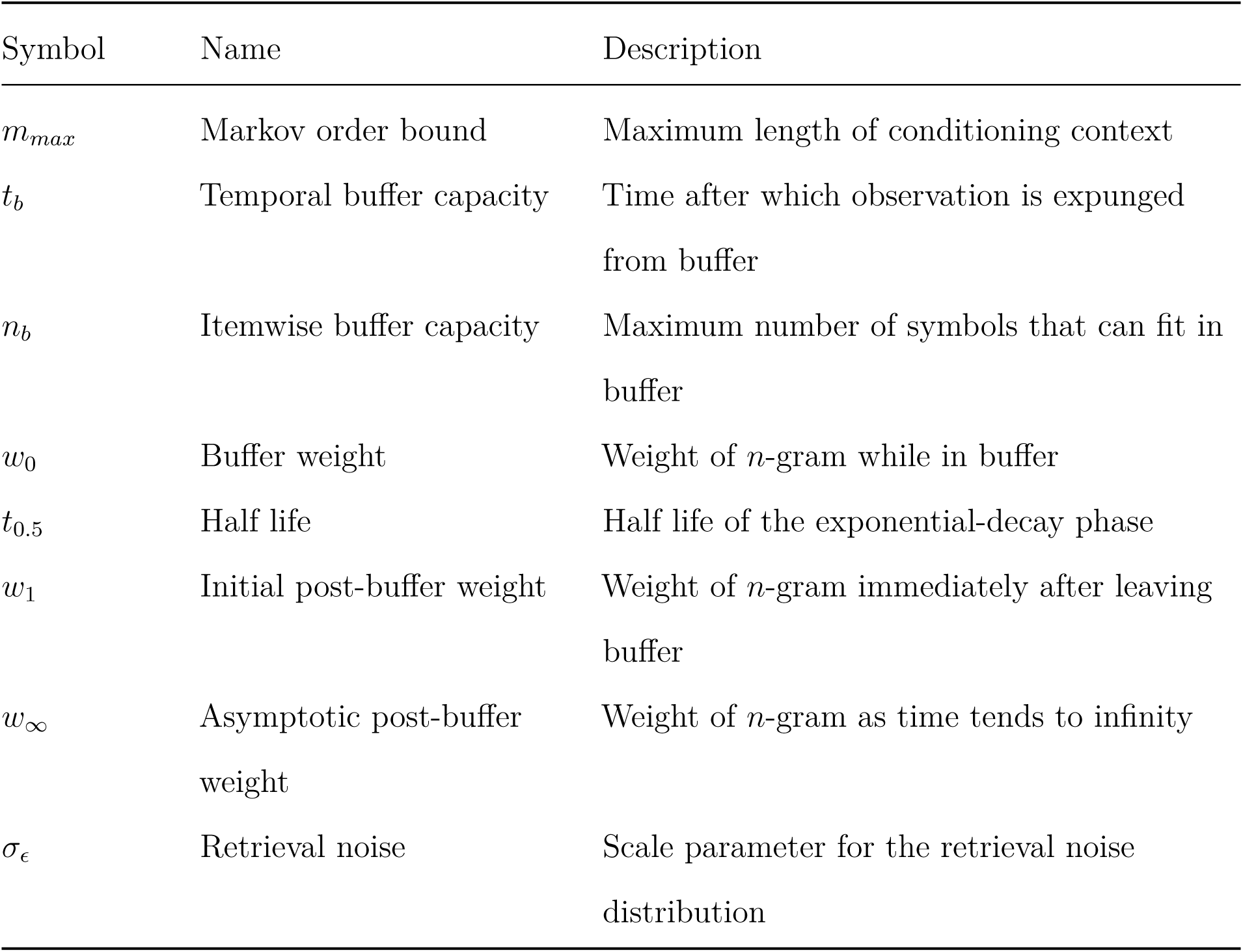
Summary of PPM-Decay hyperparameters.

### Musical corpora

#### Popular corpus

This corpus was derived from the McGill Billboard corpus of Burgoyne (2011), a dataset of popular music sampled from the Billboard “Hot 100” charts between 1958 and 1991. The sampling algorithm was designed such that the composition dates should be approximately uniformly distributed between 1958 and 1991, and such that composition popularity should be approximately uniformly distributed across the range of possible chart positions (1–100). Having sampled 1,084 compositions with this algorithm, Burgoyne (2011) had expert musicians transcribe the underlying chord sequences of these compositions. These transcriptions took a textual format, where each chord was represented as a combination of a root pitch class (e.g. “Ab”) and a chord quality (e.g. “maj”). For example, the following text represents the beginning of “Night Moves” by Bob Seger:

~~~
| Ab:maj | Ab:maj. . Gb:maj | Db:maj | Db:maj. . Gb:maj |
~~~

As is common in harmonic analyses, these transcriptions characterize chords in terms of their constituent *pitch classes*. A pitch class is an equivalence class of pitches under *octave transposition*; octave transposition means shifting a pitch by twelve semitones, which is equivalent to multiplying (or dividing) its fundamental frequency by a power of two.

This “root + chord quality” representation is intuitive for performing musicians, but it is problematic for cognitive modeling in that the chord root is a subjective music-theoretic construct. We therefore translated these textual representations into sequences of *pitch-class chords*, defined as the combination of a bass pitch class with a set of non-bass pitch classes (see Harrison & Pearce, 2020 for details). We performed this translation using the chord dictionary from the *hrep* software package (Harrison & Pearce, 2020, https://doi.org/10.5281/zenodo.2545770).

Harmonic analyses often do not systematically differentiate between one long chord and several repetitions of the same chord. In this and the following corpora we therefore collapsed consecutive repetitions of the same chord into single chords, as well as omitting all explicitly marked section repeats from the original transcriptions.

At the time of writing, only part of the Billboard corpus had been publicly released, the remainder being retained for algorithm evaluation purposes. Here we used the 739 transcriptions available at the time of writing, having removed transcriptions corresponding to duplicate compositions.

Figure 3A shows the resulting transcription for the first eight bars of “Night Moves”. The full corpus is available in the *hcorp* R package alongside the other two musical corpora used in this paper (https://doi.org/10.5281/zenodo.2545754).

#### Jazz corpus

This corpus was derived from the iRb corpus of Broze & Shanahan (2013), a dataset of lead sheets for jazz compositions as compiled from an Internet forum for jazz musicians. Broze and Shanahan converted these lead sheets into a textual representation format termed `**jazz`, which (similar to the McGill Billboard corpus) expresses each chord as a combination of a root pitch class and a chord quality, alongside its metrical duration expressed as a number. For example, the following text represents the beginning of “Thanks for the Memory” by Leo Robin:

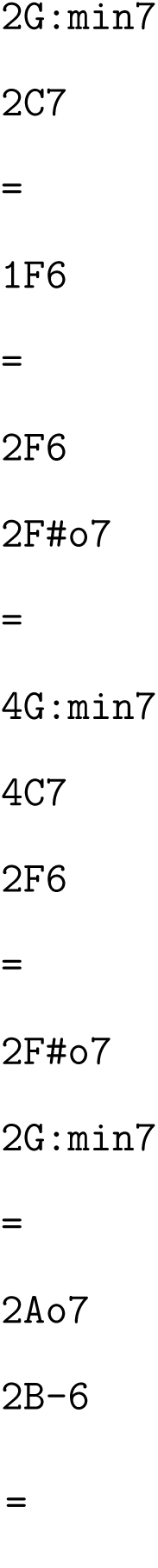

As with the popular music corpus, we translated these textual representations into sequences of pitch-class chords using the chord dictionary from the *hrep* package (Harrison & Pearce, 2020), and eliminated consecutive repetitions of the same chord. Figure 3B shows the result for the first eight bars of “Thanks for the Memory”.

#### Bach chorale corpus

This corpus was derived from the “371 chorales” dataset from the KernScores repository (Sapp, 2005). This dataset comprises four-part chorale harmonizations by J. S. Bach, as collected by his son C. P. E. Bach and his student Kirnberger, and eventually digitally encoded by Craig Sapp. The 150th chorale harmonization is omitted from Sapp’s dataset as it is not in four parts, leaving 370 chorales in total. This dataset uses the `**kern` representation scheme (Huron, 2002), designed to convey the core semantic information of traditional Western music notation. For example, the following text represents the first two bars of the chorale harmonization “Mit Fried und Freud ich fahr dahin”:

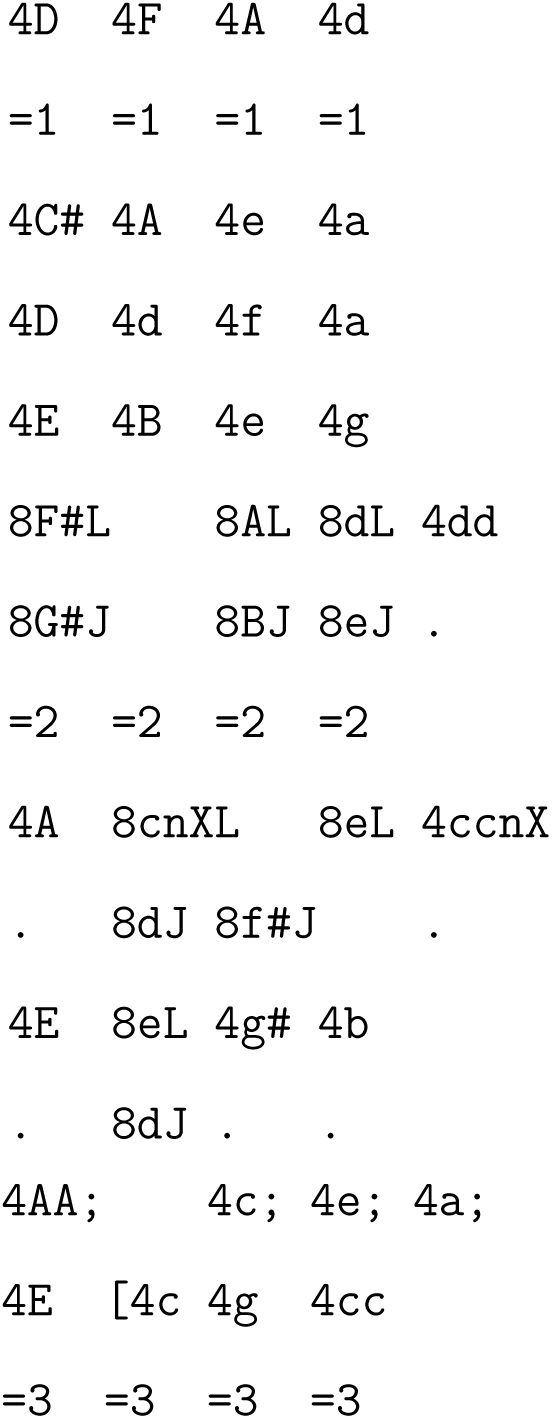

We derived chord sequences from these `**kern` representations by applying the harmonic analysis algorithm of Pardo & Birmingham (2002), which selects from a dictionary of candidate chords using a template-matching procedure. Here we used an extended version of this template dictionary, described in Table 4.

**Table 4.** The dictionary of chord templates used in constructing the Bach chorale corpus.

| Pitch classes | Label | Weight |
| --- | --- | --- |
| [0, 4, 7, 11] | maj7 | 0.2 |
| [0, 3, 7, 10] | min7 | 0.2 |
| [0, 4, 8] | aug | 0.02 |
| [0, 7] | no3 | 0.05 |
| [0, 7, 10] | min7no3 | 0.05 |
| [0, 4, 7] | maj | 0.436 |
| [0, 4, 7, 10] | dom7 | 0.219 |
| [0, 3, 7] | min | 0.194 |
| [0, 3, 6, 9] | dim7 | 0.044 |
| [0, 3, 6, 10] | hdim7 | 0.037 |
| [0, 3, 6] | dim | 0.018 |

*Note.* Each row identifies a different template.
Each template comprises a set of pitch classes, expressed relative to the chord root. Applied to a collection of pitch classes within a harmonic segment, Pardo and Birmingham's (2002) algorithm evaluates each candidate template with the respect to each of the 12 possible chord roots, and selects the template and root combination that best reflect the pitch-class content of the harmonic segment. Ties are broken using the "weight" attribute; templates with higher weights are given priority.

We computed one chord for each quarter-note beat, reflecting the standard harmonic rhythm of the Bach chorale style, and collapsed consecutive repetitions of the same chord into one chord, as before. Figure 3C shows the result for the first eight bars of the chorale harmonization “Mit Fried und Freud ich fahr dahin”.

### Behavioral experiment

#### Stimuli and procedure

Each stimulus comprised a sequence of tones, with each tone gated on and off with 5-ms raised cosine ramps. Tone frequencies were drawn from a pool of 20 values equally spaced on a logarithmic scale between 222 Hz and 2,000 Hz. Tone length was always constant within a given trial and across trials in a block. Across blocks, three different tone durations were used (25, 50 and 75 ms). Individual stimuli ranged in length between 117 and 160 tones and in duration between 3,250 and 11,025 ms.

Four stimulus types were defined: “CONT”, “STEP”, “RAND”, and “RAND-REG”. CONT and RAND trials contained no section change: CONT trials constituted one repeated tone of a given frequency, and RAND trials constituted randomly sampled tones from the full frequency pool, with the constraint that final tone counts were balanced by the end of the stimulus. STEP and RAND-REG trials each contained exactly one section change, occurring between 80 and 90 tones after sequence onset. Each section of a STEP trial comprised one repeated tone of a given frequency, with the section change constituting a change in frequency. RAND-REG trials comprised an initial random section, constructed under the same constraints as RAND trials, followed by a REG section constituting repeated iterations of a sequence of tones sampled randomly from the frequency pool without replacement. These repeating sequences comprised either 10 or 20 tones, depending on the block, with the REG section always comprising at least three repeating cycles. All stimuli were generated anew at each trial, and RAND and RAND-REG sequences occurred equiprobably.

The experimental session was delivered in 6 blocks, each containing 80 stimuli of a given tone length and alphabet size (35 RAND-REG, 35 RAND, 5 STEP, and 5 CONT), with the inter-stimulus interval jittered between 700 and 1100 ms, and with block duration ranging between 5.7 and 17.4 minutes. The order of blocks was randomized across participants. Before starting, participants were familiarized with the task with a short training session comprising six short blocks of 12 trials each, representing the same conditions as the main experiment. Stimuli were presented with the PsychToolBox in MATLAB (9.2.0, R2017a) in an acoustically shielded room and at a comfortable listening level selected by each listener.

Participants were encouraged to detect the transition as fast as possible. Correspondingly, feedback about response accuracy and speed was delivered at the end of each trial. This feedback consisted of a green circle if the response fell between the first and the second cycle of the regularity, or before 400 ms from the change of tone in the STEP condition; for slower RTs, an orange circle was displayed.

The RAND-REG trials were of primary interest for our analyses. We used the STEP trials to estimate baseline response times, computed separately for each participant within each block using correct responses only, and normalized the RAND-REG response times by subtracting these baseline response times. We excluded all RAND-REG trials where the participant responded incorrectly, and interpreted RAND and CONT trials as foils for the change-detection task.

#### Participants

We collected data from 25 paid participants (20 females; mean age 24.17, *SD* age = 3.17). Data from two participants were discarded due to overly slow reaction times on the STEP condition (mean reaction time more than three standard deviations from the mean). The research ethics committee of University College London approved the study, and written informed consent was provided by each participant.

#### Preprocessing reaction time data

We discarded 530 trials where participants responded incorrectly, and then normalized each participant’s reaction times by subtracting the mean reaction time to all correctly answered STEP trials in the same block. We then retained all RAND-REG trials where the normalized reaction times fell within two standard deviations from the mean for a given combination of participant, tone duration, and cycle length. This left 4,439 trials.

### Modeling reaction time data

We modeled participants’ reaction times using the new PPM-Decay model presented in *Model*. We modeled each trial separately, resetting the model’s memory after each trial.

We modeled participants’ change detection processes using a non-parametric change-detection algorithm that sequentially applies the Mann-Whitney test to identify changes in a time series’ location while controlling the false positive rate (Ross, 2015; Ross et al., 2011). We used the algorithm as implemented in the “cpm” R package (Ross, 2015), setting the desired false positive rate to one in 10,000, and the algorithm’s warm-up period to 20 tones.

For comparison with the participant data, we computed representative model reaction times for each condition by taking the mean reaction time over all trials where the model successfully detected a transition, excluding any trials where the model reported a transition before the effective transition (this resulted in excluding 0.41% of trials). We used R and C++ for our data analyses (R Core Team, 2017); our PPM-Decay implementation is available at https://github.com/pmcharrison/ppm and https://doi.org/10.5281/zenodo.2620414. Raw data, analysis code, and generated outputs are archived at https://doi.org/10.5281/zenodo.3603058.

